# Transcriptomic Analysis of Human Sensory Neurons in Painful Diabetic Neuropathy Reveals Inflammation and Neuronal Loss

**DOI:** 10.1101/2021.07.23.453576

**Authors:** Bradford E. Hall, Emma Macdonald, Margaret Cassidy, Sijung Yun, Matthew R. Sapio, Pradipta Ray, Megan Doty, Pranavi Nara, Michael D. Burton, Stephanie Shiers, Abhik Ray-Chaudhury, Andrew J. Mannes, Theodore J. Price, Michael J. Iadarola, Ashok B. Kulkarni

## Abstract

Pathological sensations caused by peripheral painful neuropathy occurring in Type 2 diabetes mellitus (T2DM) are often described as ‘sharp’ and ‘burning’ and are commonly spontaneous in origin. Proposed etiologies implicate dysfunction of nociceptive sensory neurons in dorsal root ganglia (DRG) induced by generation of reactive oxygen species, microvascular defects, and ongoing axonal degeneration and regeneration. To investigate the molecular mechanisms contributing to diabetic pain, DRGs were acquired postmortem from patients who had been experiencing painful diabetic peripheral neuropathy (DPN) and subjected to transcriptome analyses to identify genes contributing to pathological processes and neuropathic pain. DPN occurs in distal extremities resulting in the characteristic “glove and stocking” pattern. Accordingly, the L4 and L5 DRGs, which contain the perikarya of primary afferent neurons innervating the foot, were analyzed from five DPN patients and compared with seven controls. Transcriptome analyses identified 844 differentially expressed genes. We observed increases in levels of inflammation-associated genes from macrophages in DPN patients that may contribute to increased pain hypersensitivity and, conversely, there were frequent decreases in neuronally-related genes. The elevated inflammatory gene profile and the accompanying downregulation of multiple neuronal genes provide new insights into intraganglionic pathology and mechanisms causing neuropathic pain in DPN patients with T2DM.

## Introduction

Roughly 20% of U.S. adults report chronic pain, including about 8% of individuals who categorize their pain as severe enough to interfere with quality of life^1^^8^. Within these cases of chronic pain, about 15-25% are considered to be a caused by neuropathy, defined as pain induced by nerve damage resulting from traumatic injury, infection, auto-immune disease, or from neurotoxic substances^15^. Due to its high prevalence, the most common form of neuropathic pain arises from Type 2 diabetes mellitus (T2DM)^2^. T2DM affects roughly 25 million Americans and has an estimated future growth of ∼5% annually within the United States^35^. Approximately 50% of T2DM patients will develop some form of neuropathy, with half of those neuropathic cases considered painful (diabetic peripheral neuropathy - DPN)^31, 37^. Overall, pain is often considered the most bothersome symptom for patients with diabetic neuropathy, often reaching ratings of 6 to 10 on a 0 to 10 pain intensity scale^23, 66^.

DPN is characterized as a length-dependent neuropathy, where distal extremities such as the hands and feet are predominantly affected by pain, thereby producing a characteristic “glove and stocking” pattern^80^. The soma of the primary afferent neurons that innervate the hands and feet are known to reside in the cervical and lumbar dorsal root ganglia, respectively. Aberrant peripheral nociceptor firing is implicated in promoting pain sensitivity for individuals with DPN, since a local anesthetic nerve block prevents spontaneous pain in these patients^29^. Even though nociceptor hyperactivity is linked to heightened pain sensitivity, the only major drugs currently available that directly target the nervous system to treat pain are opioids, which can be addictive and have serious side effects^14, 81^. The DRG is a heterogeneous tissue that includes approximately 10 to 12 distinct neuronal populations that include mechanoreceptors, proprioceptors, and nociceptors of various types^46, 68, 75, 85^. The DRG neurons are also enveloped by satellite glial cells, and the axons are wrapped by myelin sheaths coming from Schwann cells. While these non-neural cells provide a protective and supportive function, the DRG exists outside the blood brain barrier, and the primary afferent neurons can be more vulnerable to the multiple metabolic disturbances occurring in the bloodstream of patients with diabetes as well as to microvascular complications^21, 28, 44^.

Research into understanding DPN, however, is complicated by the fact that diabetes is a multifactorial disorder. Chronic diabetic hyperglycemia can cause oxidative stress by both an enzymatic means through activation of aldose reductase in the polyol (sugar alcohol) pathway and with the non-enzymatic production of advanced glycation end products (AGE) via sugar addition onto proteins^9, 27^. Mitochondrial dysfunction can then ensue because of the accompanying build-up of free radicals^21^. Over time, the accumulating metabolic insult to sensory neurons overrides the capacity for regeneration and repair^11, 21, 69^. Patients with DPN generally present a heterogeneity of neuropathic symptoms with varying effects on somatic sensation^58^. The level of inflammation in DPN patients along with differences in lipid metabolism may impact neuropathic disease progression by shaping aspects like myelinated fiber density^35^. Overall, about 72% of DPN patients report that their pain worsens after the initial onset, while 12% of individuals report improvement over time and 15% report no change^23^. Additionally, some patients with distal symmetric polyneuropathy, the most common form of diabetic neuropathy, also develop autonomic neuropathy^9^.

To identify potential gene regulatory processes in humans that can contribute to DPN, we acquired DRGs from 5 DPN patients and 7 healthy controls at the time of organ donation. Gene profiles of lumbar DRGs were generated by RNA-seq analyses along with histological examinations of the DRGs to investigate potential pathophysiological changes underlying DPN. Innate inflammatory pathways appear to be upregulated in the DPN individuals, possibly in response to cell injury and/or death via alarmins or damage-associated molecular patterns (DAMPs). In contrast, we saw varying degrees of neuronal loss in four out of five of the DPN subjects that lead to coexisting declines in the expression of neuronally enriched genes. Overall, our transcriptomic analyses identify an increase in inflammation within the DRGs of individuals with DPN along with an accompanying loss in neuronally related gene expression.

## Methods

### Human Dorsal Root Ganglia Preparation

Human DRG from five individuals experiencing DPN and seven healthy controls were acquired from Anabios (San Diego, CA), all collected from the cadaveric donors with consent of the next of kin (Table 1). Both L4 and L5 DRGs were chosen as these ganglia contain the soma of the primary afferent neurons innervating the foot, particularly since DPN patients often feel pain in distal extremities. DRGs were either snap frozen or stored in RNAlater (Ambion, Austin, TX). The DPN donors include 2 males and 3 females (Age: 56.6 ± 3.9 SEM; BMI: 30.2 ± 4.1 SEM), with most reporting diabetic neuropathy for 10 or more years (1 with an incomplete medical history). The healthy controls, 4 males and 3 females (Age: 45.6 ± 3.3 SEM; BMI: 25.8 ± 2.1 SEM), were listed as either having a stroke (CVA) or head trauma. RNA was specifically extracted from one of the L5 DRGs, while a L4 DRG was used for histology.

**Table 1.** Demographics on the DRG donors used in this study. BMI = Body Mass Index, COD = Cause of Death.

| **Donor** | **Age** | **Sex** | **BMI** | **Ethnicity** | **COD** |
| --- | --- | --- | --- | --- | --- |
| DPN1 | 58 | F | 20.1 | Caucasian | CVA/ICH/Stroke |
| DPN2 | 69 | F | 28 | Caucasian | CVA/ICH/Stroke |
| DPN3 | 50 | F | 43.9 | Hispanic/Latino | Anoxia/Cardiovascular |
| DPN4 | 59 | M | 34.2 | Hispanic/Italian | Anoxia/Cardiovascular |
| DPN5 | 47 | M | 25 | Hispanic/Latino | CVA/ICH/Stroke |
| Con1 | 30 | F | 19.9 | Hispanic | CVA/Stroke |
| Con2 | 49 | F | 36.7 | Caucasian | MVA/Head Trauma |
| Con3 | 39 | F | 25 | African American | CVA/Stroke |
| Con4 | 45 | M | 25.5 | Caucasian | Anoxia |
| Con5 | 47 | M | 28.5 | Caucasian | Head Trauma |
| Con6 | 54 | M | 23.9 | Caucasian | Head Trauma/Blunt Injury |
| Con7 | 55 | M | 21.1 | Caucasian | Head Trauma |

### RNA Purification and Sequencing

Bulk RNA sequencing was performed to examine differential gene expression in multiple DRG cell types that could contribute to DPN. Additionally, bulk sequencing offers reliable statistical evaluation of the gene expression changes in the DRG that averts sparse input and provides less noise. The L5 DRG was homogenized using a Bio-Gen PRO200 (PRO Scientific, Oxford, CT) rotor-stator homogenizer with Multi-Gen 7XL probes. These were 7mm sawtooth probes that were detachable so that a clean probe could be used for each sample to avoid cross-contamination. RNA was then purified with a RNeasy Midi Kit (Qiagen, Valencia, CA) using on-column DNase-I digestion according to the manufacturer’s protocol. RNA quality was scored using a Bioanalyzer (Agilent, Santa Clara, CA) and RINs between 7-9 were obtained (Table S1), indicating some moderate degradation. Libraries were prepared using an Illumina TruSeq mRNA Sample Prep Kit (polyA+ method) (San Diego, CA) with 1 µg of total RNA and then sequenced with an Illumina NovaSeq-6000 according to the manufacturer’s protocols at the NIH Intramural Sequencing Center (Rockville, MD). Although the DRGs were collected over a roughly two-year period, library prep and sequencing were performed at the same time to avoid batch variation. Amplification was performed using 10 cycles (to minimize the risk of over-amplification), and unique dual-indexed barcode adapters were applied to each library. Libraries were pooled in an equimolar ratio and sequenced on an S4 flow cell on a NovaSeq 6000 using version 1 chemistry to achieve a minimum of 74 million 150 base read pairs. The data were processed using RTA version 3.3.4.

### RNA-seq analysis

For the human DRG RNA-seq, paired-end sequencing was performed with a read length of the 150 bp. Read quality was checked using FASTQC version 0.11.6. Trimming was performed using BBTools version 38.42 to trim 20 bp off from 5’-end, and 30 bp off from 3’-end. The alignments were performed using STAR version 2.7.2a to the hg38 reference human genome and Gencode release 27 for transcriptome annotation. Read counts per gene per sample were quantified using HTSeq version 0.9.1. Table S1 shows alignment statistics and the total number of reads mapped to genes per sample. A list of differentially expressed genes (DEGs) between the DPN and the healthy samples was generated using DESEQ2 version 1.24.0^49^. Sex of the donor was treated as a covariate in the design when building the generalized linear model for DESEQ2 analysis, hence, removing unwanted variation. With adjusted p-value cutoff of 0.05 by Benjamin Hochberg’s False Discovery Rate (FDR), differential expression returned 411 genes up-regulated genes and 433 genes down-regulated genes in the diabetic neuropathy condition compared to the healthy controls (Tables S2, S3, and S4). Principal component analyses were performed before and after the covariate controlling of sex of the donor, which show improved separation of DPN vs. Control along PC1 axis (Figure S1).

### Pathway and Data Analysis

All 844 dysregulated genes were entered into Ingenuity Pathway Analysis (IPA) (Qiagen, Valencia, CA) to identify enriched pathways and potential upstream regulators. Because of the robust inflammatory signal detected in the DRG of the DPN donors, DEGs were split between upregulated and downregulated genes to determine which pathways were affected by decreased gene expression levels. The STRING Database (Version 11.0) (https://string-db.org/) was used to look for interactions between significantly differentially expressed genes. STRING recognized only 593 of the 844 differentially expressed genes. Enriched Gene Ontology (AMIGO)^72^ terms identified using STRING mostly related to inflammatory processes and included immune response GO:0006955 (89 genes with a false discovery rate of 8.17e-06) and cytokine-mediated signaling pathway GO:0019221 (47 genes; false discovery rate of 5.09e-05). Neuronal genes were recognized with STRING as well, particularly synapse genes (GO:0045202). Enrichment analysis was also done using ToppGene Suite (https://toppgene.cchmc.org), which identified dysregulated CD markers (HGNC group ID 471 - https://www.genenames.org/). Heatmaps were generated using Seaborn in Python, version 3.8.3. All heatmaps display the Z Score of Log2 Transformed Normalized DESEQ2 Gene counts. In order to further evaluate DRG compositional changes resulting from DPN, the DEGs from the human donors were compared with bulk RNA sequencing of rat DRG and sciatic nerve, which serve as surrogates for peripheral neurons and Schwann cells respectively^64^. A heatmap was created to display the counts of the DPN human sample sequencing counts for the overlapping gene list, while also labeling the genes enriched in the neuronal or Schwann cell group. The z score by gene of the log2 corrected counts are displayed. The density plot was created with the log2 corrected fold change for all genes that overlap from the enriched for neuronal or Schwann cell gene sets (not just the differentially expressed). This shows the trend towards up or downregulation amongst the entire gene enrichment list. In order to calculate the likelihood of the bias of the enriched datasets towards up or downregulation in the human DRG dataset, a hypergeometric test was performed for each enrichment gene list. A scatterplot of the DEGs was additionally generated using GraphPad Prism (Version 8.0.2) in order to compare log2 fold change to base mean of normalized DESeq2 counts.

### Histology and Immunohistochemistry

With signs of inflammation and neuronal loss appearing in our transcriptome analysis, a L4 DRG from 5 control and 5 DPN donors was subsequently sent to Histoserv (Germantown, MD) for paraffin embedding and sectioning. Frozen L4 DRGs were temporarily held at −20°C for cutting purposes, sliced in half, and immediately placed in fixative (10% buffered formalin). After paraffin embedding, 6μm sections were cut, and slides were stained with hematoxylin and eosin along with luxol fast blue (data not shown) for histological analysis. Immunohistochemistry was also performed by Histoserv (Germantown, MD) for CD3 (AB17143 CD3 antibody [F7.2.38], Abcam, Cambridge, MA), CD20 (AB9475 CD20 antibody [L26]), and CD68 (AB783, CD68 antibody [PG-M1]). Slides were deparaffinized in xylene and graded alcohols. The slides were then subject to either citrate-based (CD20), or tris/EDTA-based (CD3 and CD68) antigen retrieval. Then, sections were blocked and incubated with primary antibody (CD20, 1:100 dilution; CD3, 1:100 dilution; CD68, 1:500 dilution). Immunodetection of the CD markers was performed using a polymer-based reagent. After detection, slides were dehydrated and then cover-slipped in mounting medium. Stained slides were scanned on an Aperio AT2 whole slide imaging system (Leica Biosystems, Buffalo Grove, IL) and images were evaluated using FIJI to quantify average pixel density normalized over the number of hematoxylin stained nuclei identified by the program for each image^17^. Paired t-tests were used to determine differences in CD marker expression between DPN and control DRG.

### RNAscope Analysis

Formalin fixed paraffin embedded (FFPE) sections from the L4 DRG were used for in situ hybridization (5 controls and 5 DPN donors). Three neuronal genes (TRPV1, SLC17A7, and PRDM12) were labelled using RNAscope® technology, which has been previously described^76^. Using custom software, paired double-Z oligonucleotide probes were designed against target RNAs, as follows: Hs- TRPV1-C3, cat no. 415388-C3, NM_080706.3, 20 pairs, 1922 – 2905; Hs-SLC17A7-C2, cat no. 415618-C2, NM_020309.3, 20 pairs, nt 1311 - 2570; and Hs-PRDM12, cat no. 559968, NM_008509.2, 20 pairs, nt 117 – 1789. TRPV1 and SLC17A7 probes were stained for five and four different images (respectively), while PRDM12 was stained for one image for each sample. The RNAscope LS Multiplex Fluorescent Reagent Kit (Advanced Cell Diagnostics, Newark CA) was used on FFPE sections according to manufacturer’s instructions, with target retrieval using ER2 buffer at 95°C for 15 minutes, and protease III at 40°C for 15 minutes as pre-treatment. Each subject was quality controlled for RNA integrity using an RNAscope Human 4-plex Positive Control Probe (Advanced Cell Diagnostics, Newark CA). Fluorescent images were acquired with a 3D Histech Scanner. For each scanned image, three neuronally dense areas were selected for analysis. Three reviewers naïve to tissue sample condition evaluated all neurons in these areas for expression of TRPV1, SLC17A7, and PRDM12. Cochran-Mantel-Haenszel tests for repeated tests of independence for two probes (TRPV1 and SLC71A7), and Fisher’s Exact Test was performed for one probe (PRDM12) to determine significance.

## Results

### Comparative Transcriptomic Analysis of DPN versus Healthy Controls

To gain molecular insight into causes of diabetic neuropathic pain in patients, we recovered DRGs from consenting organ donors experiencing diabetic painful neuropathy (DPN, Table 1). DPN is a length-dependent neuropathy, affecting the distal extremities such as the fingers and toes, so L4 and L5 DRGs were recovered as these ganglia contain the soma of the primary afferent neurons innervating the foot. DRGs were recovered from 5 DPN donors along with 7 controls. Total RNA was extracted from the L5 DRGs, and mRNAs were enriched (polyA+ method). Bulk RNA-seq was conducted to holistically analyze expression changes in the DRG that can occur with DPN, as the DRG is comprised of not only sensory neurons, but also Schwann cells, satellite glial cells, fibroblasts, perivascular cells, and resident and infiltrating immune cells^64^. Sequencing data from the DRGs was aligned to the human genome and differential gene expression analysis was performed using DESEQ2^49^. After examination of the RNA-seq data, principal component analysis was used to examine how the donors would cluster. Initially, the DRG donors were clustered by sex, but, after treating sex as a covariate, we had a clear separation between DPN versus healthy controls as the principal division amongst the donors (Figure S1). Overall, between the two groups, we were able to derive 844 differentially expressed genes (p-value < 0.05), with 411 genes significantly upregulated in donors with DPN versus 433 genes significantly downregulated (Figure 1, Table S2).

**Figure 1.**
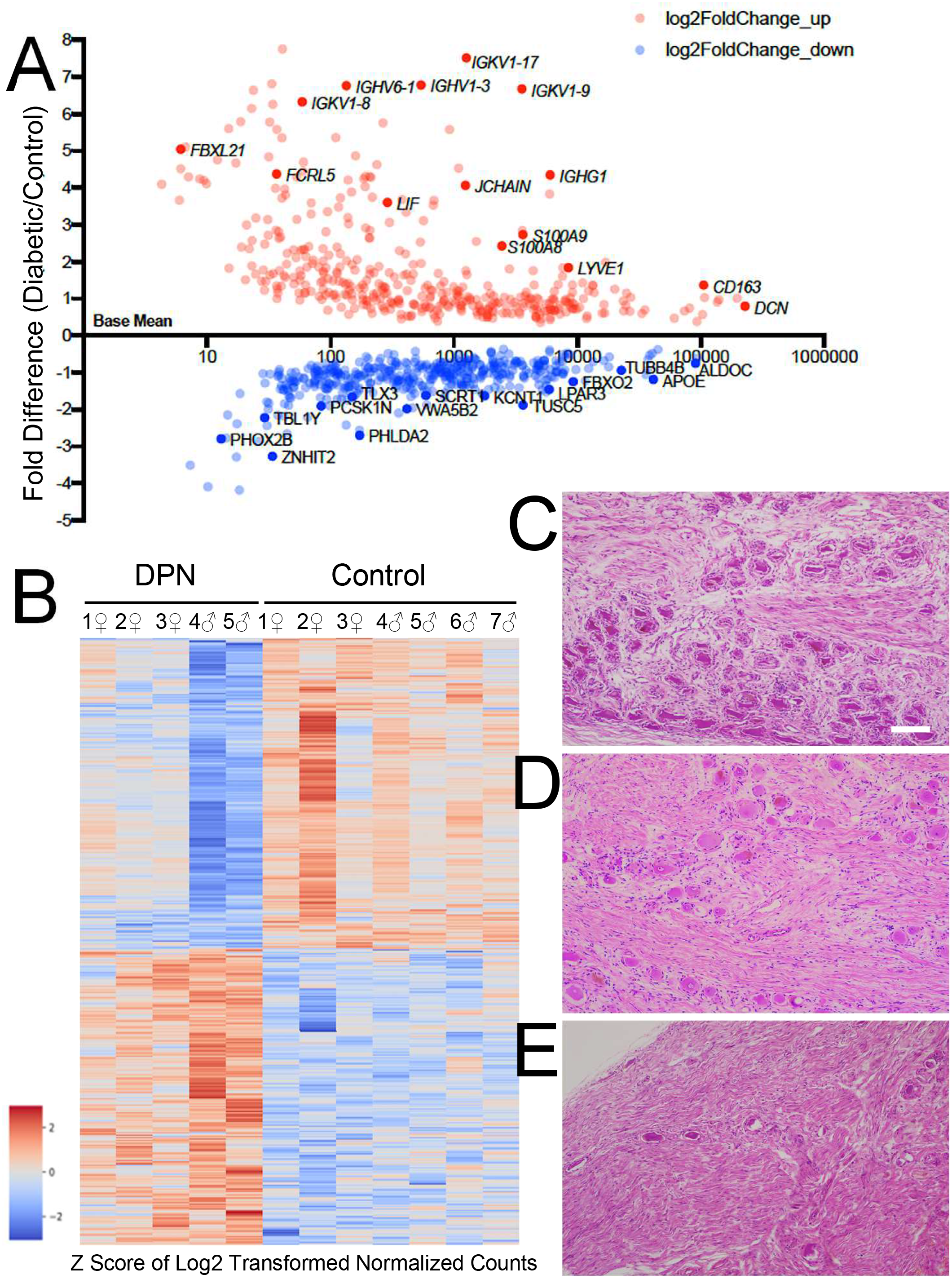
DPN Transcriptome: A) Scatterplot of all DEGs where dysregulated genes with high log2 fold change per base mean of normalized DESeq2 counts are highlighted. Immunoglobulins and other inflammatory related genes are upregulated while some genes involved in neurogenesis like the transcription factors SCRT1 and TLX3 were downregulated in the diabetic individuals. B) Heatmap of all 844 DEGs. C-E) H and E images of the L4 DRG, scale bar = 500 µm; C) Normal dorsal root ganglion with adequate density of ganglion cells surrounded by satellite cells, D) DPN with moderate loss of neuronal ganglion cells with replacement reactive fibrosis, and E) DPN with severe loss of neuronal ganglion cells with replacement fibrosis (hematoxylin and eosin stains at 40X magnification).

As seen in Figure 1A, we saw that the upregulated genes detected in our transcriptomic analysis include immunoglobulins in conjunction with other inflammatory response genes, while, in the same samples, neuronally related genes were essentially downregulated. We analyzed our list of DEGs with Ingenuity Pathway Analysis (IPA) to determine which biological processes were most affected by diabetic neuropathy. We divided our data set between upregulated and downregulated genes. Analysis of the upregulated genes emphasized the increased inflammatory responses within the DRG of the DPN donors, where the top canonical pathways involve communication between immune cells and diapedesis (Figure S2A). IPA also predicted activation of TNF-α, IL-6, and IL-1β cellular signaling pathways, where these key immunoregulators are known to be associated with diabetes-induced metabolic inflammation^24, 67^. In contrast, with the downregulated genes, we detected canonical pathway changes associated with synaptogenesis signaling (Figure S2B). Patients with diabetic neuropathy often develop sensory loss over time even while still experiencing pain and our pathway analysis similarly appears to show a transcriptomic loss of neuronally related gene expression in the DRG of the DPN individuals^22^.

We next histologically examined the L4 DRGs from the donors in order to see if there were pathological changes that would support our findings in the RNA-seq data (Figure 1C-E). Of the 5 control DRGs available for histopathological examination, one was mostly normal (Con1), three had mild loss of myelinated axons (Con 3, 6, and 7), while one individual (Con4), with Lennox-Gastaut Syndrome, had mild to moderate neuronal ganglionic cell loss. In contrast to the controls, a broad span of neuropathology was seen for the DPN individuals (Figure 1D and E). One of the female donors (DPN3) was apparently normal, while another female (DPN2) showed mild to moderate neuronal ganglionic cell loss, and a third female (DPN1) had moderate neuronal loss with scattered degenerating neurons. Of the two DPN males, DPN4 showed moderate loss of neuronal ganglionic cells with scattered degenerating neurons and loss of myelinated fibers while DPN5 had moderate to significant neuronal loss with scattered nodules. Greenbaum et al.^26^ also reported a varied extent of pathology in DRGs from diabetic neuropathy patients experiencing pain that ranged from minor degenerative changes to loss of neurons with the appearance of nodules of Nageotte in one female patient.

A heatmap of DEGs in Figure 1B depicts not only a clear contrast between the DPN patients versus the controls, but also illustrates a range of gene expression changes occurring within the DPN DRGs as well, probably due to differences in disease pathology and progression. The heatmap shows that there is evident downregulation in the expression of several genes in the two DPN males that likely stems from advanced neuronal loss. Two of the three female DPN subjects also have a loss of neurons, but don’t exhibit the same degree of gene expression change as the two males. Nonetheless, the gene expression decreases in these female DPN donors probably still result from neurodegeneration within the DRG. Of note, patients with diabetic neuropathy often show a reduction in distal epidermal nerve fiber density^44^, and this axonal degeneration can also impact neuronal gene expression in the DRG neurons. Sex is known to differentially affect both diabetic onset and pain intensity^1, 5^, but, with our limited sample size, we cannot confirm if there is a true sex-linked difference causing the distinct male and female gene expression changes in figure 1B or if this pattern is just a consequence of variation in disease progression. Despite this range in neuropathic symptomology, our transcriptomics is still able to identify common trends amongst the individuals with DPN.

### Transcriptional Regulators, Ion Channels, Kinases, and G-protein coupled receptors

In general, pain hypersensitivity often occurs downstream of inflammation, where inflammatory mediators are known to activate transcription factors and trigger kinase activity in nociceptors. Therefore, we specifically examined the expression changes in transcriptional regulators and kinases in our data set to search for differences between the DPN individuals and the controls that might impact neuropathic pain (Figure 2). For transcriptional regulators (Figure 2A), we saw an overall upregulation of known T-cell induced transcriptional activators in the DPN individuals including Nuclear Factor Interleukin 3 Regulated Protein (*NFIL3*) and Interferon Gamma Inducible Protein 16 (*IFI16*). We additionally saw that BCL6 Transcription Repressor (*BCL6*) and its corepressor *BCOR* are increased in the DPN donors, which may be indicative of inflammatory cell infiltration. The Class II Major Histocompatibility Complex (MHC) Transactivator *CIITA* is increased as well in the DPN donors, which can act as a macrophage antigen presentation gene. In contrast, for neurons, we observed downregulation in several transcriptional regulators that are enriched in nociceptors. Transcription factors like *ISL2*, *TLX3*, and *PRDM12*, for example, were all significantly downregulated in the DPN donors^71^. Other downregulated neuronal transcription factors include *NHLH1*, *SCRT1*, *SCRT2*, and *SIX4*. Interestingly, *INSM2*, a gene expressed within sensory neurons, was downregulated in the DPN subjects too, where this transcriptional repressor is additionally known to influence insulin secretion and glucose tolerance^77^.

**Figure 2.**
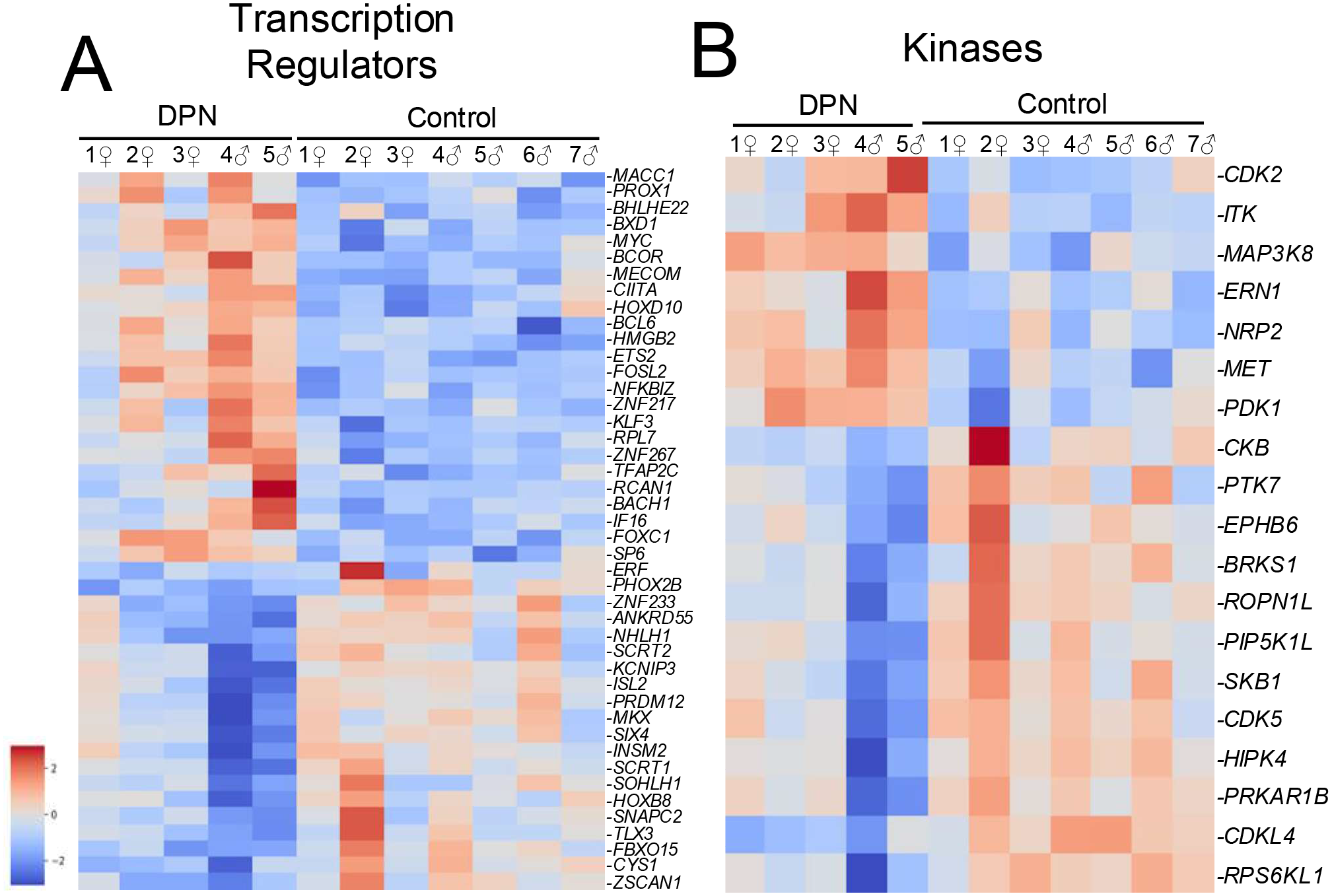
Heatmaps of dysregulated genes belonging to functional protein classes associated with pain signaling. Pictured are heatmaps of A) transcriptional regulators and B) kinases. Inflammation can cause gene expression changes in nociceptors through downstream intracellular signaling cascades that ultimately activate various transcription factors. Inflammatory signaling can additionally stimulate protein kinases that can then modulate pain transducing ion channels to induce peripheral sensitization.

When examining for changes in kinase expression, inflammation-related genes again were upregulated in the DPN individuals, including *ITK* and *MAP3K8*, both of which are again commonly expressed by T-cells (Figure 2B). In the same dataset, three creatine kinases were surprisingly downregulated in the DPN individuals, including *CKB*, *CKMT1A*, and *CKMT1B*, all of which could impact energy utilization in neurons. *PRKAR1B*, a subunit of the PKA holoenzyme, is also downregulated. The expression of *CDK2*, a key cell cycle kinase, is conversely increased in the DPN individuals and could broadly suggest either local immune cell activation, tissue repair, or neuronal apoptosis. The HGF receptor *MET* is also upregulated within the DPN individuals, which can alternatively indicate either inflammation or possible axonal regeneration^51^. Lastly, *NRP2*, a kinase linked with axonal regeneration during Wallerian degeneration^3^, was upregulated in the individuals with DPN.

Nociceptor firing can be modulated by ion channel activity and through GPCR signaling, so, next, we examined the expression of genes that encode for these two protein classes within our list of differentially expressed genes (Figure 3). With few exceptions, most altered ion channels were downregulated (Figure 3A). Notably, almost half of the downregulated ion channels in the DPN individuals were neuronally expressed potassium channels, including the potassium voltage-gated channels *KCNH2* and *KCNQ2* (both predominantly expressed in Aβ-fibers), the inwardly rectifying channel *KCNJ11*, the calcium-activated channel *KCNN1*, and the sodium-activated channel *KCNT1*. In addition to these potassium channels, two potassium channel-interacting proteins, *KCNIP2* and Calsenilin (*KCNIP3*) were additionally downregulated within our dysregulated gene list as well, where both of these calcium binding proteins are known to affect neuronal excitability by modifying A-type potassium channels. The downregulation of these potassium channels could potentially lead to either more neuronal hyperexcitability or could just be indicative of the neuronal loss that was seen histologically. Besides potassium channels, *ASIC3* (acid sensing ion channel 3), a prominent pain transducing ion channel expressed in C-fibers, was downregulated in the DPN donors too, possibly in conjunction with the loss of other neuronal markers. However, *GLRA3*, a glycine receptor that may contribute to prostaglandin E2 related pain hypersensitivity, is a neuronally associated gene that is surprisingly upregulated within the DPN subjects^30^.

**Figure 3.**
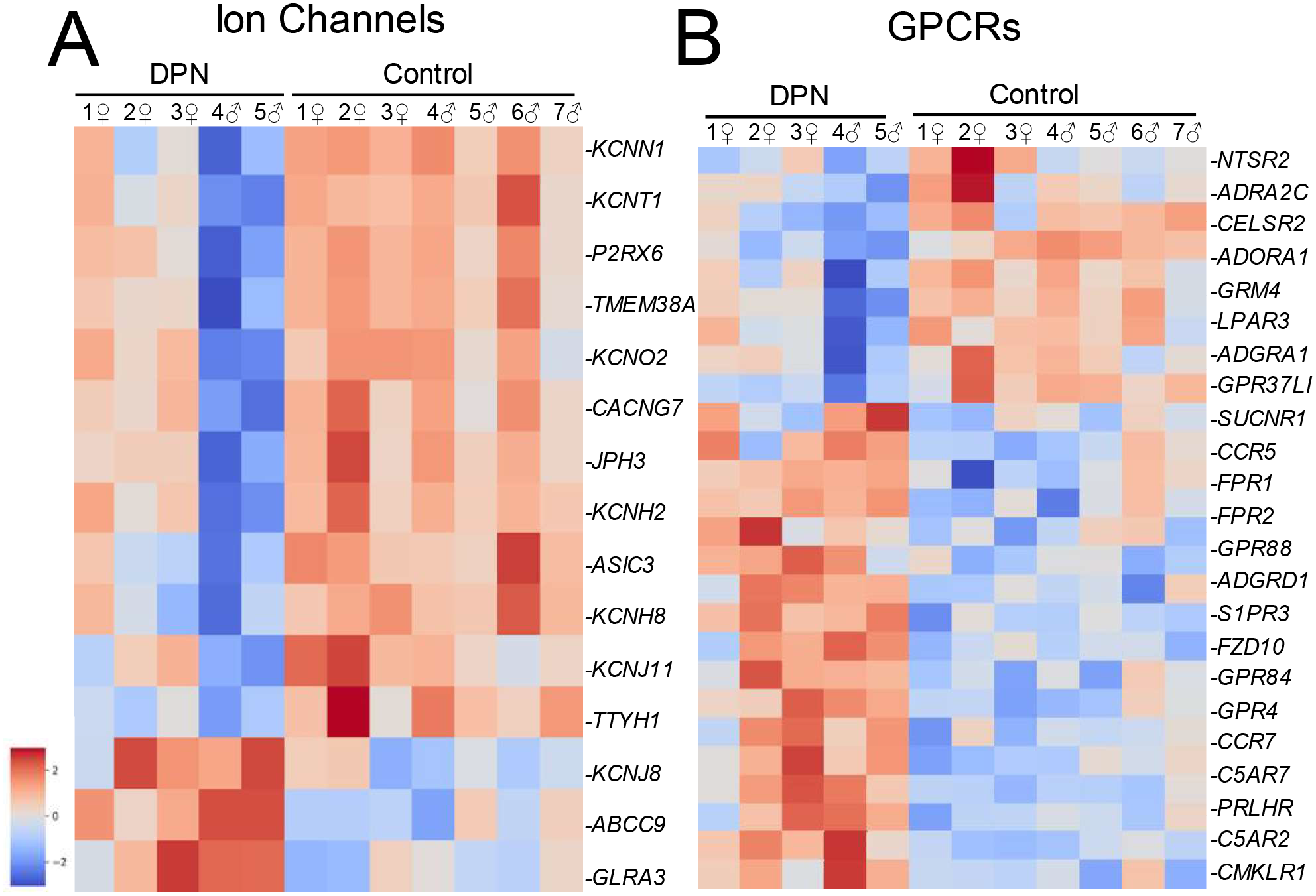
Heatmaps of genes dysregulated in DPN individuals that encode for A) ion channels and B) G protein coupled receptors. Ion channels and GPCR signaling can affect nociceptor firing. Transcription factors can alter the expression of key ion channels and GPCRs that can subsequently promote nociceptor hyperactivity. Kinases can modify the activity of ion channels and GPCRs through post-translational phosphorylation to then regulate the sensitivity to noxious stimuli in nociceptors.

Amongst GPCRs, inflammation-related signaling pathways were once more significantly upregulated as neuronal modulatory genes were downregulated in the DPN individuals (Figure 3B). *C5AR1* and *C5AR2*, receptors for the proinflammatory chemoattractant complement C5a, were both upregulated in the DRGs from the DPN donors. The formyl peptide receptors *FPR1* and *FPR2* were upregulated in the DPN individuals as well, both of which are mutually involved in the detection of cellular damage. In terms of pain signaling, the DPN donors have increased expression of sphingosine 1-phosphate receptor *S1PR3*, a GPCR that is known to promote mechanical pain partially via closure of KCNQ2, which coincidentally is downregulated in the DPN donors^32^. DPN subjects also showed decreased expression of receptors for adenosine (*ADORA1*), norepinephrine (*ADRA2C*), and dynorphin (*OPRK1*), all of which possess antinociceptive qualities. Downregulation of these receptors can either cause increased pain hypersensitivity or again be applicable to neuronal loss, yet proenkephalin (*PENK*), the pro-form ligand for μ opioid receptor, is interestingly upregulated in our list of genes, possibly in response to diabetes-induced metabolic stress. This contrasts with animal models of axon injury, where the *Pdyn* transcript that encodes for the dynorphin opioid prepropeptide is upregulated^65^. With our bulk sequencing, however, the full implication for some of these example gene expression pattern differences cannot be fully understood without further analysis to identify which cell types have been affected.

### Downregulation of neuronal marker genes

Along with the pathological neuronal loss seen histologically, our transcriptome analysis also shows a corresponding decrease of additional neuronally-related genes in the DPN donors (Figure 4). Of particular note is the significant downregulation of the NeuN gene (*RBFOX3*), a well-established neuronal marker. Synuclein Gamma (*SNCG*), a gene highly expressed in the peripheral nervous system, shows decreased expression in the DPN donors as well. *LPAR3*, which is highly enriched in a subpopulation of TRPV1+ neurons, and PVALB, which labels DRG myelinated proprioceptors, are both additionally downregulated in the DPN individuals and supports the idea that diabetic neuropathy ultimately impacts multiple sensory neuron types^68^. Additionally, the DPN individuals also show downregulation of *NAT8L*, a key enzyme known to catalyze the formation of the metabolite N-acetylaspartate (NAA), which is important for neuronal health.

**Figure 4.**
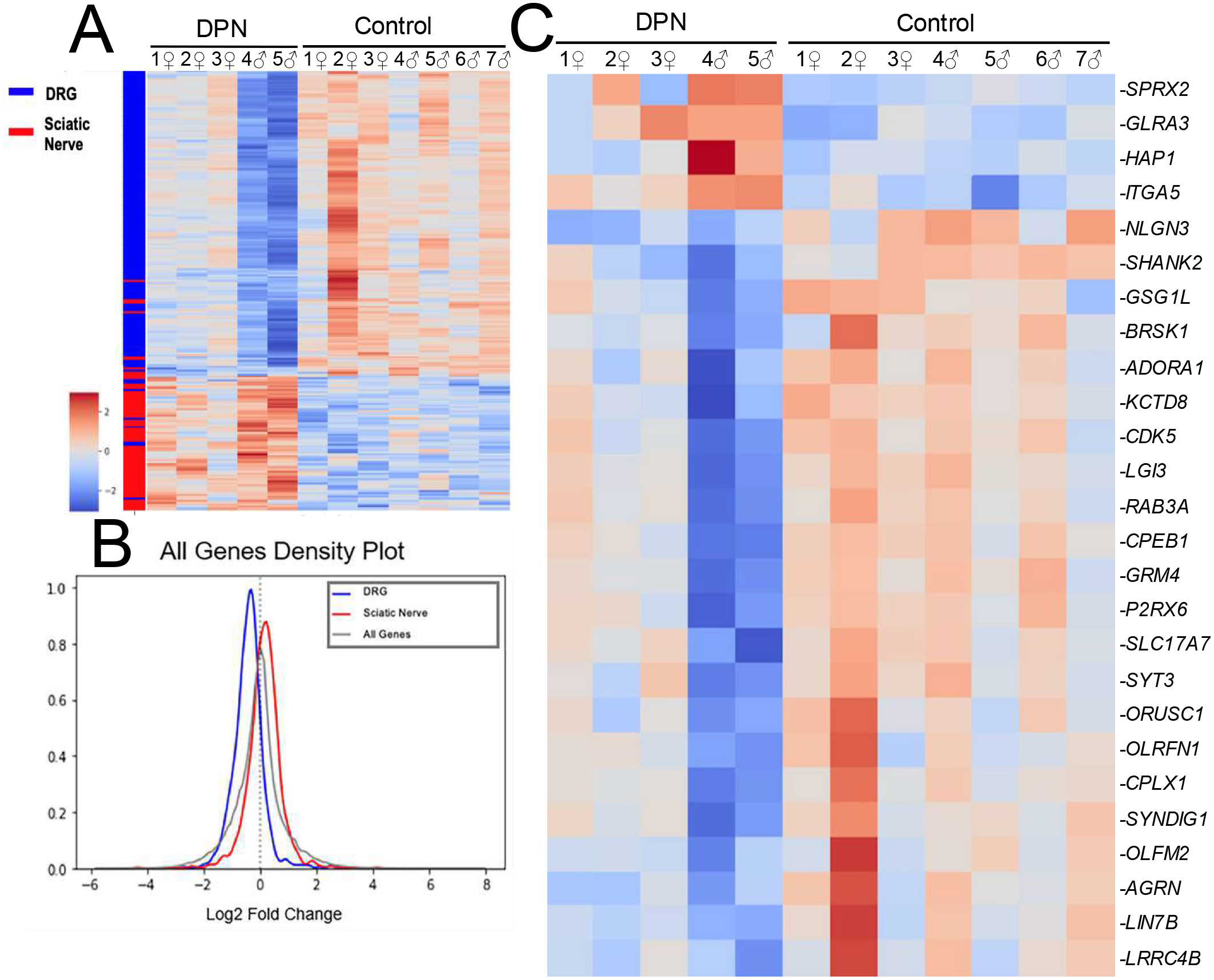
Downregulation of neuronal cell type specific genes: A) Our transcriptomic list of DEGs was compared with a database of rat homologs enriched in the DRG (which contains the soma of sensory neurons) versus those enriched in the sciatic nerve (which predominantly consist of Schwann cells that support descending neuronal axons). B) Neuronally related genes (DRG) were essentially downregulated in the DPN individuals, particularly in the two male DPN donors. Consequently, genes more related to Schwann cell character (sciatic nerve) predominate in the data and are configured as appearing upregulated due to this loss in neuronal gene density. C) Of the neuronally specific genes identified in our list of DEGs, enrichment analysis characterized 41 genes as related to synaptic function.

With the transcriptomic loss of neuronal genes in our list of DEGs, we decided to compare our RNA-seq data with a dataset of genes enriched in the rat DRG and sciatic nerve^64^ (Figure 4A and B). While the DRG contains both neurons and Schwann cells, the DRG is enriched in neurons while the sciatic nerve is predominately comprised of Schwann cells, which allows for identification of genes enriched in these two cell types. In general, most neuronally enriched genes are downregulated compared to Schwann cell genes. Dysregulated neuronal genes were then further characterized according to cellular function. Of the genes categorized as a structural component of a neuron, about 66% were associated with the synapse while 39% are considered as part of the neuronal cell body (Figure S3A). In particular, genes linked to synaptic vesicle exocytosis and the synaptic vesicle cycle are downregulated in the DPN individuals including *CPLX1*, *SYT3*, and *RAB3A* (Figure 4C). Key neuronal kinases that affect neurotransmitter release including *BRSK1* and *CDK5* were downregulated also (Figure S3B). In addition, single-pass membrane proteins important in synapse assembly are downregulated such as the *SDK2* (sidekick cell adhesion molecule 2) and *LRFN1* (leucine rich repeat and fibronectin type III domain containing 1). As seen in Figure 4C, the DPN individuals additionally have decreased expression of *CPEB1*, a regulatory protein that controls the translation of mRNA that is localized to the synapse. Surprisingly, some notable neuronal genes are upregulated in the differential gene expression analysis despite the general neuronal loss seen in the DPN individuals. Ephexin1 (*NGEF*), for example, is upregulated, possibly in response to neuronal injury^62^. Ephexin is primarily known to participate in axonal guidance but can switch to inhibit axonal regeneration following activation of the ephrin receptor EphA4^63^.

Lastly, to investigate the neuronal loss further, in situ hybridization was performed on sections of L4 DRGs in order to visualize and quantify the expression of three neuronally associated genes: *TRPV1*, a known pain transducer highly expressed in nociceptive Aδ and C-fibers^54^; *SLC17A7*, a glutamate transporter that labels large diameter sensory neurons; and *PRDM12*, a key transcriptional regulator of nociceptors during development (*SLC17A7* and *PRDM12* were both significantly downregulated in our transcriptome analysis). Consistent with the histological report, *TRPV1*, *SLC17A7*, and *PRDM12* were expressed in the DPN individuals in a lower proportion of DRG neurons than in control DRG neurons (Figure 5). Expression of all three genes was significantly decreased at the p < 0.0001 level. Essentially, the loss of normal neuronal markers demonstrated in transcriptomic results was evident in our ISH fluorescence labeling including decreased neuronal density in the DRG along with decreased neuronal gene expression in DRG neurons. Despite the loss of nociceptors, there are residual primary afferent neurons that could still trigger spontaneous pain^29^.

**Figure 5.**
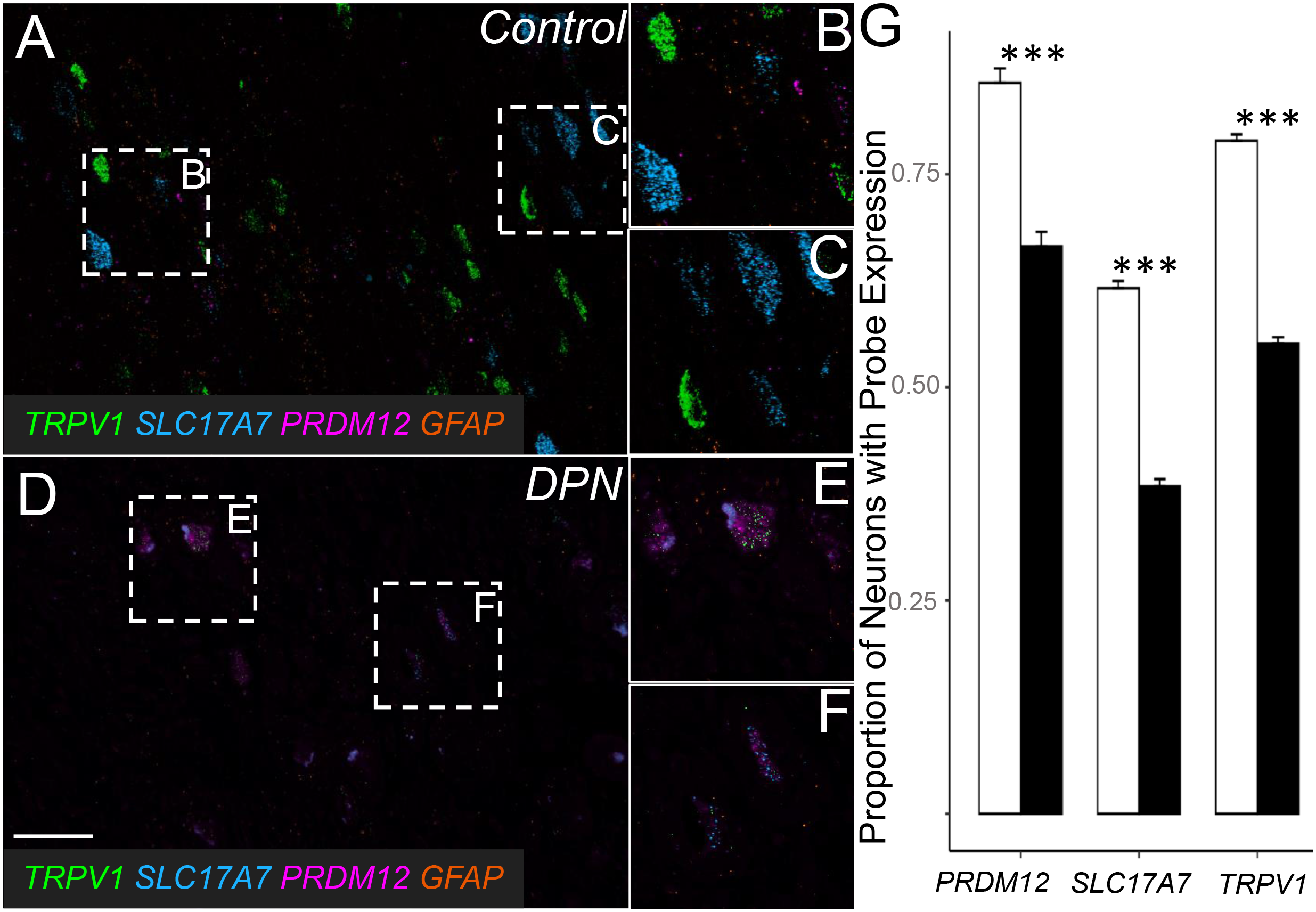
Representative images and comparison of RNAscope data: RNAscope data confirms a loss of neuronally related gene as noted in RNA-seq results by demonstrating a decrease in the proportion of neurons expressing each neuronal probe. A) DRG of healthy individuals displayed B) small diameter TRPV1 positive neurons and C) larger diameter SLC17A7 positive neurons. D) DRG of DPN individuals had decreased proportions of neurons expressing all three probes, while still containing expected neuronal subtypes such as E) small TRPV1 positive and F) larger SLC17A7 positive neurons. Scale bar = 100 µm. G) Differences in proportion of neurons expressing each RNAscope between control and DPN tissue were significant at the ***p < 0.0001 level, according to the Cochran-Mantel-Haenszel Test (TRPV1 and SLC17A7), and Fisher’s Exact Test (PRDM12). Each probe was significantly downregulated, evidence of a loss of neuronally related gene expression in DPN DRG. Data is shown graphically as mean and standard error. The downregulated expression of these three neuronal markers coincides with the neuronal loss seen histologically. Although dedifferentiation could cause a loss of TRPV1+ and SLC17A7+ neurons, most neurons were still labelled with either of these two markers.

### Increased inflammation in the DRG due to DPN

Overall, our transcriptome analysis shows a downregulation in neuronal marker genes while, conversely, displaying an increased immune cell signature, probably as a result of inflammatory cell infiltration. We see indications of both macrophage and T-cell recruitment to the DRG in DPN patients, which is similarly seen in rodent models of neuropathic pain^20^. We surprisingly also see the induction of an antibody immune response in the DPN individuals. (Table S5). Essentially, immunoglobulins account for both ∼17% of all significantly upregulated genes (71 of the 411) and mark 39 of the top 100 most significantly altered genes, all of which may reflect an aberrant B cell polyclonal activation that has been reported to be circulating in the blood of patients with diabetes^84^. Transcriptome analysis, however, cannot determine if the immunoglobulin transcripts are targeted towards neurons or supporting glia. Besides immunoglobulins, we detected upregulation in numerous genes related to an overall immune response (Figure 6A). Upregulated inflammatory genes are seen across all DPN individuals, which coincides with the increased metabolic inflammation that is known to develop in patients with diabetes^24, 67^. We see, for example, that S100A8/A9 are both increased in the DPN donors, where elevated circulating levels of these proteins can act as a biomarker for diabetic induced inflammation^78^. S100A8/A9 are mainly produced by neutrophils (53) and macrophages but may also be expressed by Schwann cells post-injury (12). Further analysis of the dysregulated inflammatory responses revealed that aspects of innate, humoral, and T-cell immune activity are all upregulated in the DRGs of the DPN individuals (Figure S4).

**Figure 6.**
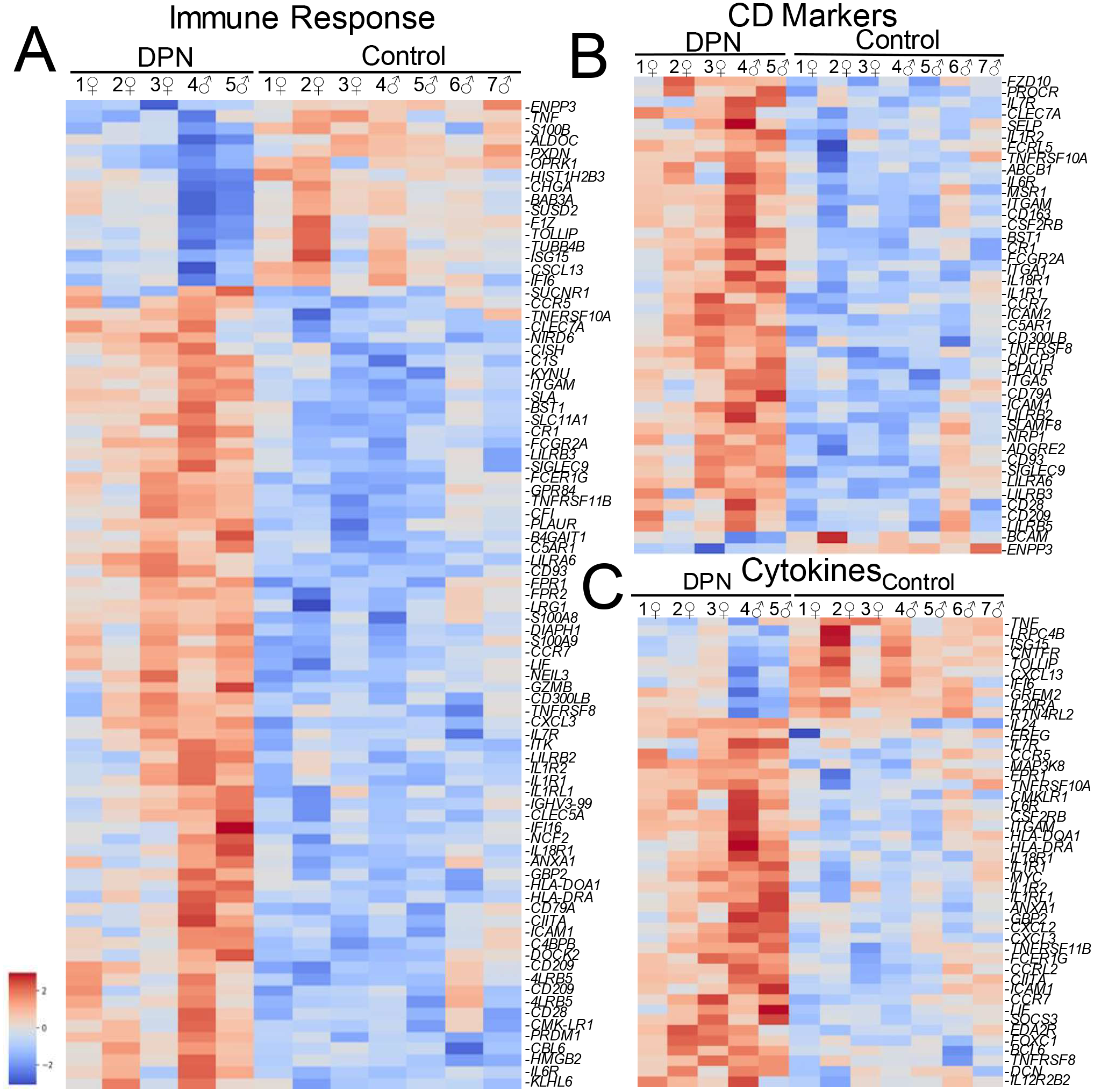
Dysregulated immune response in the DPN individuals: A) Gene enrichment analysis identified 89 genes associated with immune responses in our list 844 DEGs. As shown in the heatmap, most genes were upregulated in the DRG of the DPN subjects while only a few inflammatory genes were downregulated. B) 47 genes were further characterized as involved specifically in cytokine-mediated signaling. C) 44 genes were considered CD markers.

We next searched for differentially expressed genes that encode both for cluster of differentiation (CD) cell surface markers and cytokines to further characterize the intra-DRG inflammatory process. In figure 6B, *TNFRSF8* (CD30), a marker of activated T and B cells was upregulated in the DPN individuals. Additional markers of B-cell activation include upregulation of *CD93*, which is required for maintained antibody secretion^13^, and *KLHL6*, which is involved in B-lymphocyte antigen receptor signaling^45^. The macrophage marker *PLAUR*^10^ is also upregulated in the DPN subjects together with an accompanying increase in *ICAM1* (CD54). Upregulated ICAM1 expression may indicate the recruitment and extravasation of neutrophils into the DRG, which is consistent with the presence of S100A8 and A9^53^. Accompanying the pro-inflammatory response within the DRG of the DPN subjects, M2 macrophage scavenger receptors including *MSR1* (CD204) and *CD163* were both upregulated in the DPN individuals as well, where M2 macrophages are generally known to be involved in the phagocytic clearance of cell debris and the resolution of inflammation, perhaps pointing to a mechanism as to how neuronal debris may be cleared.

In terms of cytokine signaling (Figure 6C), several members of the interleukin 1 receptor family were upregulated including *IL1R1*, *IL18R*, and *IL1RL1* (IL33R). Also, of interest is the increased expression of *IL6R* in conjunction with elevated *IL1R1* as high circulating levels of both IL1β together with IL-6 are predictive of developing T2DM^67^. In contrast, the DPN donors show increased expression of *IL10*, an anti-inflammatory cytokine that is often considered protective against neuroimmune diseases^47^. In terms of chemokine signaling, inflammation in the DRG of the DPN donors appeared to be accompanied by increased expression of the leukocyte chemoattractant ligands *CXCL2* and *CXCL3*, otherwise known as macrophage inflammatory protein 2-alpha and beta respectively, along with *CCRL2*, which is upregulated during monocyte infiltration and macrophage differentiation. Lastly, the DPN subjects exhibit upregulation of the chemokine receptor *CCR5*. Although expressed on various immune cell types, the increased expression of *CCR5* in the DPN donors could indicate a shift towards a M1 proinflammatory macrophage status^43^.

The presence of inflammatory response genes in our DRG transcriptomic dataset suggests both an activation of resident immune cells along with leukocyte infiltration into the DRG of the DPN donors. To better understand the nature of the increased immune response in DPN DRG (i.e., cell-mediated, humoral, or innate), immunohistochemistry was performed for CD3, a T-cell marker, CD20, a B-cell marker, and CD68, a macrophage marker. The staining intensity for CD3 (Figure 7A and B) and CD20 (Figure 7D and E) were both intriguingly higher in the DPN subjects versus the controls but were not statistically significantly so, even despite the upregulation of B and T-cell markers in our transcriptome analysis. However, increased staining for CD68 (Figure 7G and H) was statistically significant within the DRG of those with DPN, demonstrating that the upregulated inflammatory markers captured in the transcriptomics are likely linked to innate immune responses, potentially stemming from diabetes related neuronal pathology. Our immunohistochemistry staining suggests that macrophage-driven inflammation may have a more predominant impact on the DRG of the DPN donors.

**Figure 7.**
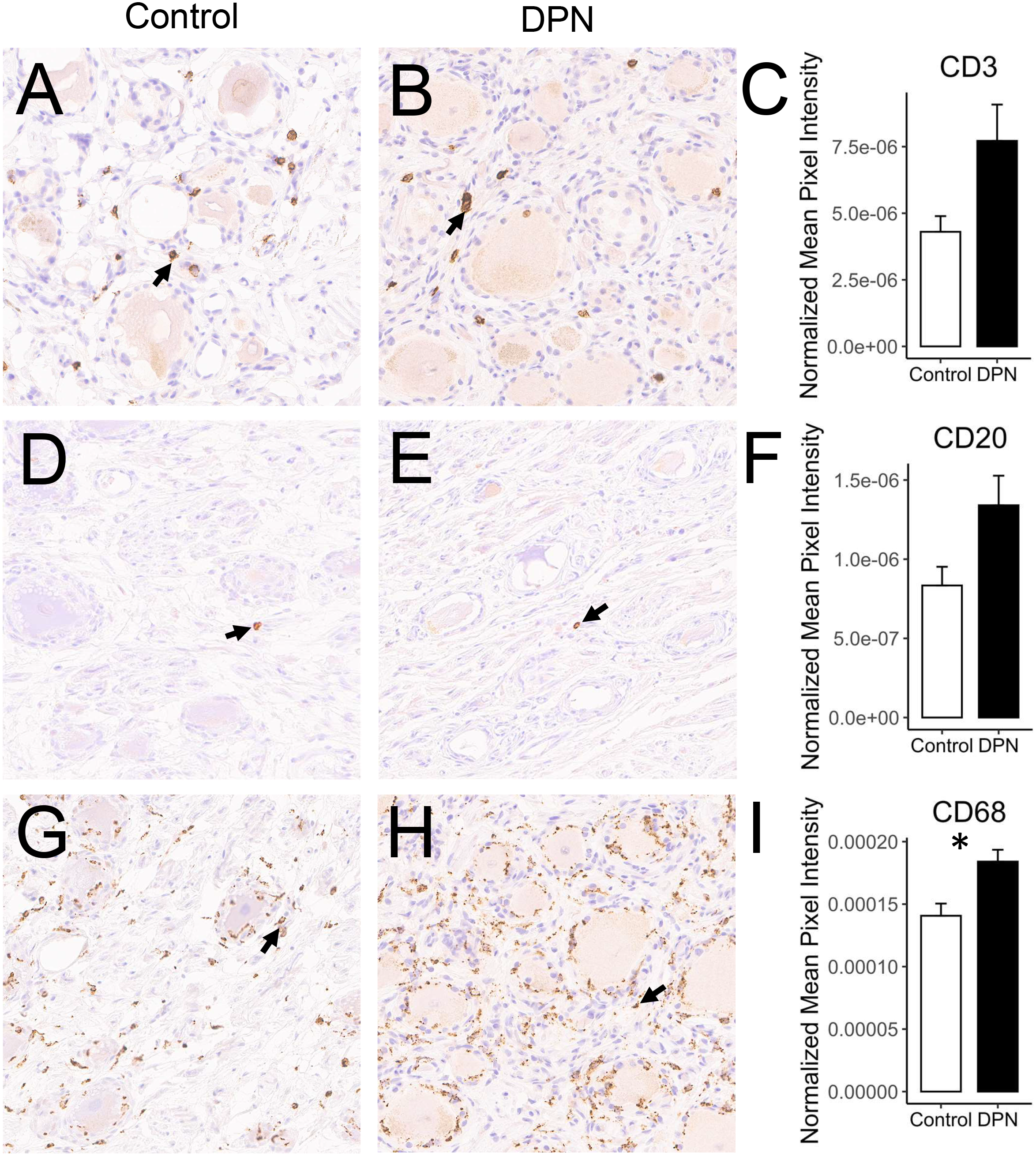
Three CD markers were chosen to characterize the immune related responses observed in the transcriptome: A and B) CD3, a T-cell marker, D and E) CD20, a B-cell marker, G and H) and CD68, a macrophage marker. Scale bar = 100 µm. Stained images were scanned and processed using FIJI to quantify average pixel density normalized over the number of hematoxylin stained nuclei for each image (83). Paired t-tests were performed to compare average expression of CD3, CD20, and CD68 between DPN (B, E, and H) and control DRG (A, D, and G). Average differences in expression were higher for all three markers in the DRG of the DPN individuals, but only CD68 was significant (C, F and I) (*p < 0.05 level, one-tailed paired t-test), suggesting the activation primarily of innate macrophage driven immune responses.

## Discussion

T2DM can cause axonal loss, demyelination, and pain in about a quarter of patients^31, 37, 74^. The metabolic disorder resulting from T2DM can subsequently cause hyperexcitability in sensory C-fiber neurons that, in turn, may promote neurodegeneration in this chronic illness^37^. Neuronal injury can eventually also affect neighboring “spared” nerve fibers^20^. As with other forms of neuropathic pain, there are no effective treatments for DPN^15^. In general, about 39% of patients with DPN do not receive any treatment for pain^19^, while, of those who are treated, only one-third achieve at least a 50% therapeutic reduction in pain symptoms^38^. Existing therapeutics also cannot reverse neuropathy once it has developed^86^. Like other types of neuropathic pain, treatment with traditional chronic pain medications such as nonsteroidal anti-inflammatory drugs or opiates is often ineffective in alleviating DPN, and most clinical drug trials have failed to produce any significant therapeutic outcomes^2, 8, 21^.

In order to better understand the underlying pathology associated with painful neuropathy, we performed transcriptomic analysis using DRGs recovered from organ donors who died with DPN. The DRG is a heterogenous tissue, so bulk RNA-seq was conducted to identify possible contributing factors such as microangiopathy, Schwann cell dysfunction, or aberrant macrophage activation. Only a few pain studies have used human DRGs that contain the neuronal soma^25, 59^. Our study uniquely uses RNA-Seq analysis to compare DRG gene expression differences in healthy controls versus a cohort of subjects that all share the same chronic pain condition. We were also able to make clear connections between the gene expression changes uncovered through RNAseq with the histopathology seen in the DRG, particularly in terms of neuronal loss and activation of T-cell and macrophage populations.

Overall, inflammation can be an important factor following neuronal injury, as macrophages are needed to clear cellular debris. Sustained inflammation, however, can be detrimental to neuronal viability^52^. Our transcriptome analysis appears to indicate aberrant inflammatory processes are occurring in the DRGs of the DPN donors (Figure 6A). Low-grade inflammation (“metaflammation”) is generally known to be linked to diabetes^24, 34^. The DRGs collected for our transcriptomic analysis were obtained late in the disease course, so we cannot determine if inflammation was a major factor in the development of painful neuropathy or if the inflammatory signature seen in our RNA seq data occurred later in response to nerve damage. Nonetheless, monocytes isolated from peripheral blood of T2DM patients have been shown to exhibit higher levels of proinflammatory markers including IL-6, IL-1α, TNF-α, and ICAM1, which supports the notion of low-grade inflammation and infiltration in the pathogenesis of diabetes^24^. Although *TNF* (TNF-α) was surprisingly downregulated, the expression levels of *ICAM1* along with the interleukin receptors *IL1R1* and *IL6R* were all elevated in the DPN donors in our study (Figure 6A)^50^. IL-1β and IL-6 are key regulators of inflammatory reactions^52^ and together are predictive for the risk of developing T2DM^67^. IL-1β and IL-6 are both known to stimulate nociceptor firing and are additionally both reported to be upregulated in sural nerve biopsies of patients with progressive diabetic neuropathy^6, 7, 35^. Chronic hyperglycemia arising from T2DM can cause the activation of macrophages that then often proceed to infiltrate into adipose tissue, the liver, and other tissues. We likewise see increased staining for *CD68* in the DRG of the DPN donors (Figures 7G and H) as well as the upregulated expression of macrophage markers including *PLAUR*, *MSR1*, and *CD163* in our transcriptomic data set (Figure 6B). The inflammation seen in the DPN donors suggests a reaction to signaling by alarmins that are released in response to cell death^82^. The signature of a macrophage driven immune response was similarly detected in a transcriptomic analysis of the db/db mouse model of type 2 diabetes^33^. Macrophages are known as key regulators of neuropathic pain and depletion of macrophages in rodent models can attenuate hyperalgesia^42, 57^. Besides expressing pro-inflammatory cytokines, macrophages can also contribute to neuropathic pain by recruiting neutrophils to the site of injured neurons, particularly through the release of *CXCL2* (macrophage inflammatory protein 2-alpha), a chemokine that is interestingly upregulated in our data set as well^41^. The resident macrophages within the DRG itself can play a significant role in promoting neuropathic pain besides the macrophages in the periphery that are reacting to metabolic inflammation^83^. Nonetheless, pro-healing M2 macrophages are probably present in the DRG of the DPN donors, too, as suggested by upregulation of the macrophage scavenger receptors *MSR1* and *CD163* and the increased expression of *IL10*.

DRG neurons and their axonal projections lie outside the blood brain barrier and are less protected than neurons within the CNS, but these primary afferent neurons have, in general, a known capacity for axonal regeneration following nerve damage^52^. In our transcriptomic results, some upregulated neuronal genes in the DPN individuals indicate possible attempts at neuronal regeneration including *NRP1* and *NRP2*^3^. However, axonal degeneration probably overrides regeneration, which may account for the predominant transcriptomic loss of neuronal genes within the DRGs of those exhibiting DPN^11, 21, 69^. In nerve transection mouse models of neuropathic pain, there is a downregulation within the sensory ganglionic neurons of neuron-specific genes, similar to our findings, yet there is also a concurrent induction of injury-induced genes like *Atf3*, which was not observed in our transcriptomic analysis of the DRG of DPN individuals, possibly because of the different nature of the neuropathy or the clinical versus experimental time-course^55, 61^. In addition, while the decrease of neuron-specific genes seen in our transcriptomic analysis could partly be explained by the reduced expression of key transcriptional regulators in the DPN patients^71^, more likely, the downregulation in neuronal markers probably just reflects the histopathological findings of neuronal loss in some of the DPN subjects (Figure 4 and 5).

Patients with DPN often report experiencing spontaneous pain that is described as sharp and burning and our transcriptomic analysis identified several genes reported to be linked to nociceptive signaling (Tables S6 and 7)^29^. Some aspects of this heightened pain sensitivity may be immune related, as we detected upregulation of multiple inflammatory markers. Interestingly, however, key antinociceptive receptors such as *ADORA1*, *ADRA2C*, and *ORPK1* show decreased expression in the DPN donors (Figure 3B). Additionally, several potassium channels are downregulated that are known to hyperpolarize a neuron’s membrane potential to inhibit neuronal firing. A decrease in these channels in the DPN individuals may contribute to spontaneous pain sensations (Figure 3A). Downregulated expression of *KCNQ2* (K_v_7.2), *KCNN1* (SK1), and *KCNT1* (SLACK) in particular are associated with increased sensitivity to pain^79^. The decreased expression of these potassium channels and antinociceptive receptors may either promote nociceptor hypersensitivity or, again, be indicative of neuronal loss. A few neuronally related genes are intriguingly upregulated though, including, for example, *S1PR3* and *GLRA3*, both of which are involved in promoting pain hypersensitivity. Attempts at axonal regeneration may also play a role in causing painful diabetic neuropathy versus non-painful neuropathy^11^. Axonal outgrowth may require the reactivation of some genes associated with neuronal development, so we provide a list of such upregulated genes in table S8. Surprisingly in our transcriptomic analysis, many of the voltage gated channels, ion channels, and neuropeptides that are involved in pain hypersensitivity were not upregulated in association with DPN even though increased expression of these proteins has been hypothesized to contribute to peripheral sensitization, particularly in rodent models of neuropathic pain^16, 69^.

Pain is commonly experienced early during DPN, even at a prediabetic stage^56, 60^, but, in our study, the DRGs of the DPN subjects were collected late in the course of diabetic neuropathy, after ongoing neuronal insults had occurred. So, our dataset does not reflect for the initial etiological factors in diabetes that result in pain. Our RNA-seq data basically focuses on the gene expression changes within the peripheral neurons of the DRG, but central sensitization also plays an important role in causing chronic pain in the DPN patients^70^. For example, nerve injury can cause 2^nd^ order nociceptive interneurons in the dorsal horn to undergo dendritic restructuring that leads to wider receptive fields^40^. Loss of neuronal inputs to the spinal dorsal horn may also shift inhibitory/excitatory circuit balance leading to enhanced pain signaling through remaining nociceptive inputs to the dorsal horn. MRI studies additionally suggest that possible spinal cord atrophy, abnormal thalamic function, and changes in higher brain areas may contribute to DPN as well^70^.

With our transcriptomic study, we see an increase in proinflammatory responses with a concurrent loss in neuronally related genes within the DRG of the DPN individuals. Patients with diabetic neuropathy ultimately lose feeling in their legs and feet as subjects who show higher sensory loss often paradoxically report more severe neuropathic pain^22, 58, 69, 73^. Our transcriptomic data concur with these reports of sensory loss as there is a decrease in neuronally related genes. Signs of potential neurotoxic processes can be seen in the transcriptomic analyses, for example, with upregulation of the ER stress sensor *ERN1* (Ire1-α) along with *TXNIP*^48, 60^. This is also consistent with previous studies showing activation of the ER stress pathway in peripheral nerves in rodent diabetic models and stimulation of ER stress in nociceptors by the diabetic metabolite methylglyoxal^4, 36^. The neuronal loss in the DPN subjects could alternatively result from Golgi degeneration, as reported in a rat model of diabetes^39^. In conclusion, our unique approach to transcriptomic profiling of DRGs obtained from diabetic neuropathy organ donors provides new insights into the pathology and mechanisms of DPN and raises new questions that can be explored in future experimental studies.

## Acknowledgements

We would like to thank Dr. Kenneth Yamada for critical reading of the manuscript. We also want to thank Drs. Janice Lee, Jacki Mays, Eva Mezey, Michaela Prochazkova, Wanjun Chen, Anand Swaroop, Robert Morell, and Daniel Martin along with Mr. Matthew Brooks, Mr. Gilberto Carmona, and Mrs. Anita Terse for their helpful discussions. We would also like to thank Anabios Inc. for procuring the human DRGs, and the NIH Intramural Sequencing Core (NISC) for running the RNAseq analysis. This research was supported by the Intramural Research Program of the NIDCR (ABK) and Clinical Center (AJM), NIH, and the University of Texas at Dallas and NIH grant NS111929 (TJP).

---

**Supplemental Figure 1.**
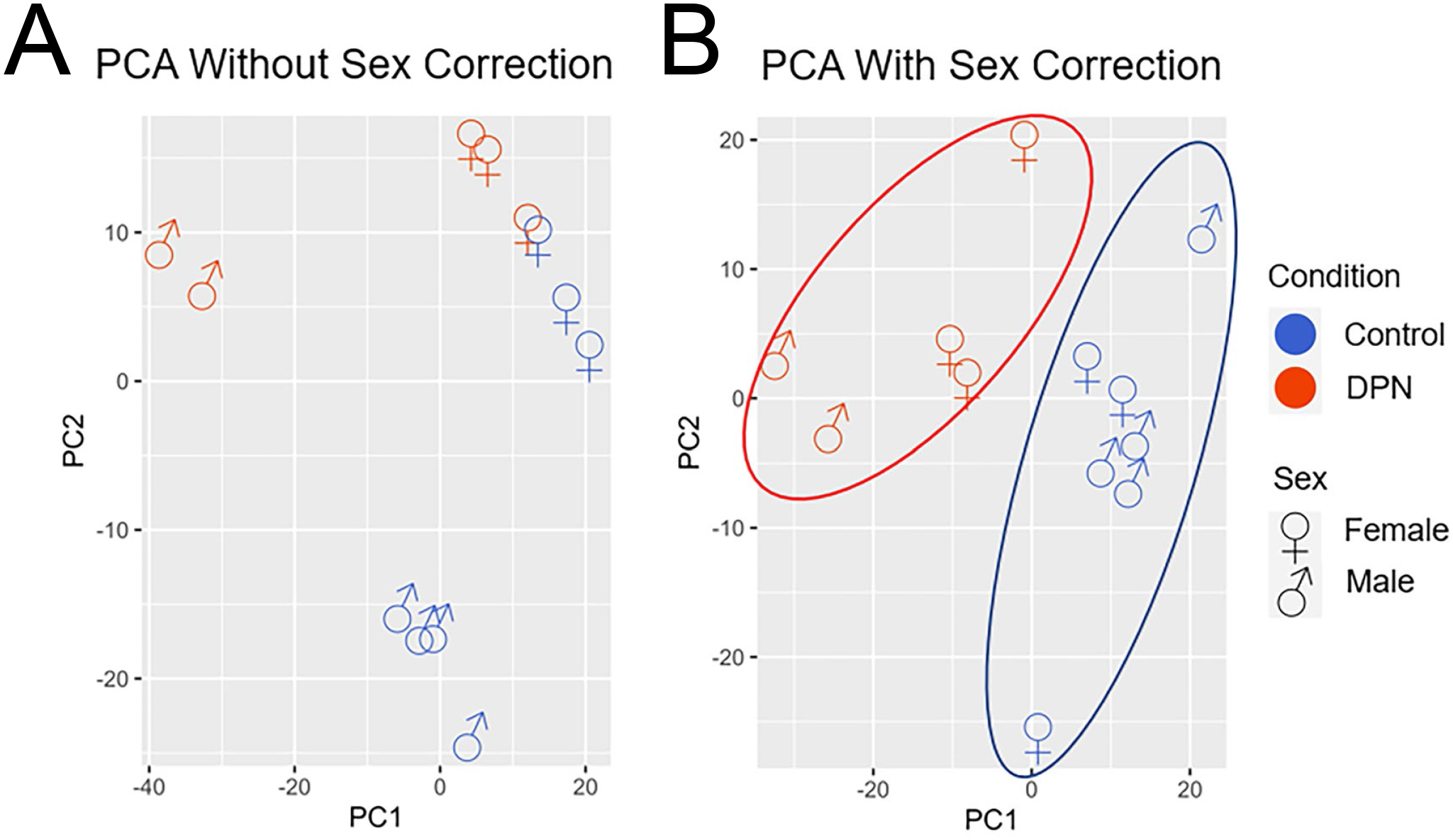
Principal component analysis (PCA) of human DPN transcriptome data: PCA plot A) before and B) after including sex as a covariate as part of the DESEQ2 analysis. PC1 is the first principal component direction where the most variance is occurring, and PC2 is the second most one that is orthogonal to PC1. With sex as a covariate, the DPN donors and the controls then separate into two independent groups.

**Supplemental Figure 2.**
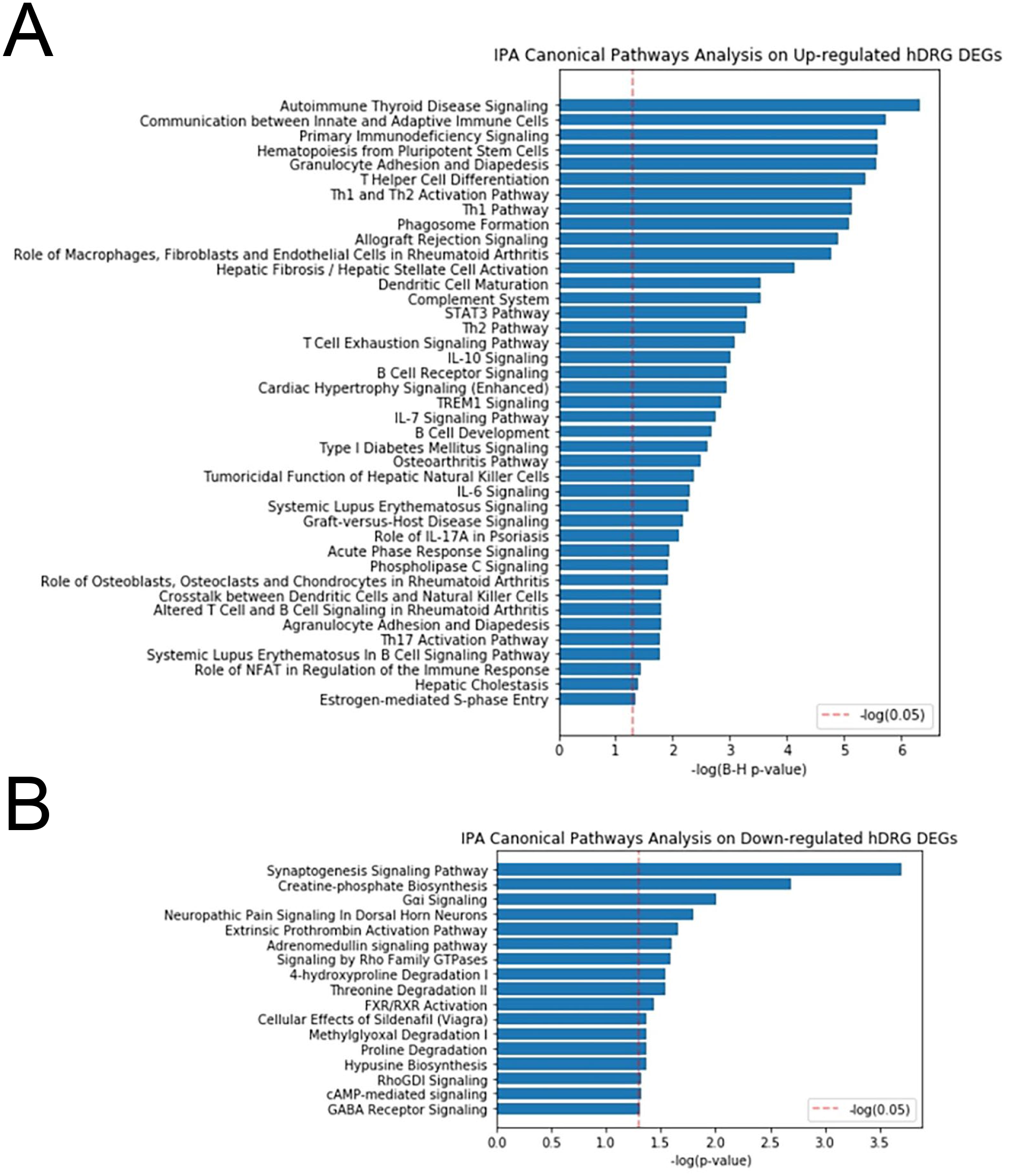
Ingenuity pathway analysis (IPA) of transcriptomic data: The IPA report using all 844 dysregulated genes predominantly centered on the immunological functions occurring in the DRG of the DPN individuals. Thereby, further assessment of the DEGs was separated between A) the upregulated largely inflammatory gene responses and B) the downregulated gene expression changes, where synaptogenesis was appears to be affected by decreases in gene expression.

**Supplemental Figure 3.**
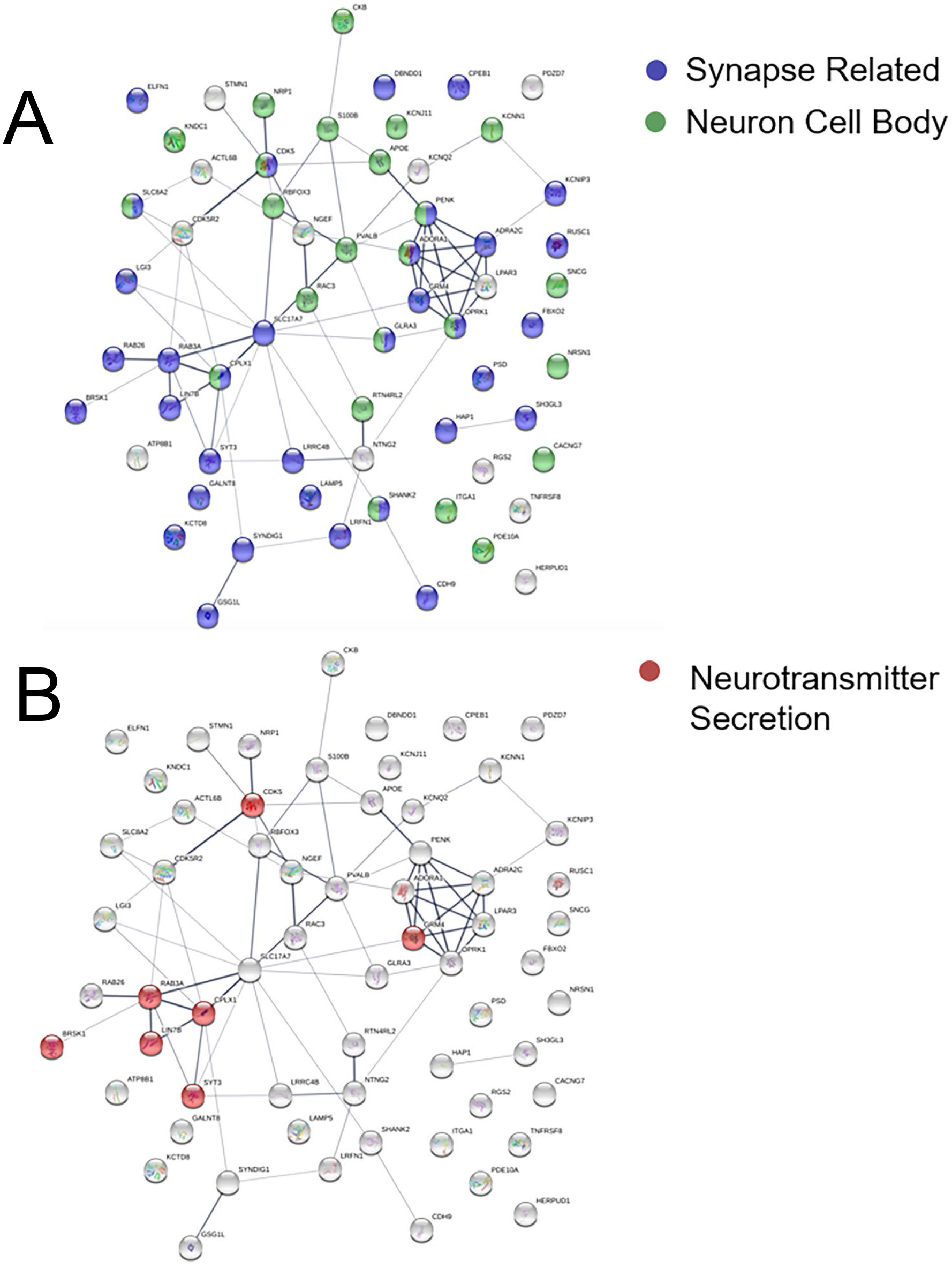
Interaction network of neuronally related genes: 62 genes out of our DEG list were considered to perform as a cellular component of a neuron (GO:0097458). Most genes were downregulated (n=51) while a few were upregulated (n=11) in the DPN donors. To further determine how these dysregulated genes might impact neuronal function, all 62 gene were separately evaluated using STRING (https://string-db.org/) for additional enrichment analysis. A) About 66% of the neuronal genes (n=41) were synaptically related (blue - GO:0045202), while 39% (n=24) were associated with the neuron cell body (green - GO:0043025). B). In terms of biological function, a few genes in red were qualified as being associated with neurotransmitter secretion (GO:0007269). Kuleshov MV, Jones MR, Rouillard AD, Fernandez NF, Duan Q, Wang Z, Koplev S, Jenkins SL, Jagodnik KM, Lachmann A, McDermott MG, Monteiro CD, Gundersen GW, Ma’ayan A. Enrichr: a comprehensive gene set enrichment analysis web server 2016 update. Nucleic Acids Research. 2016; gkw377

**Supplemental Figure 4.**
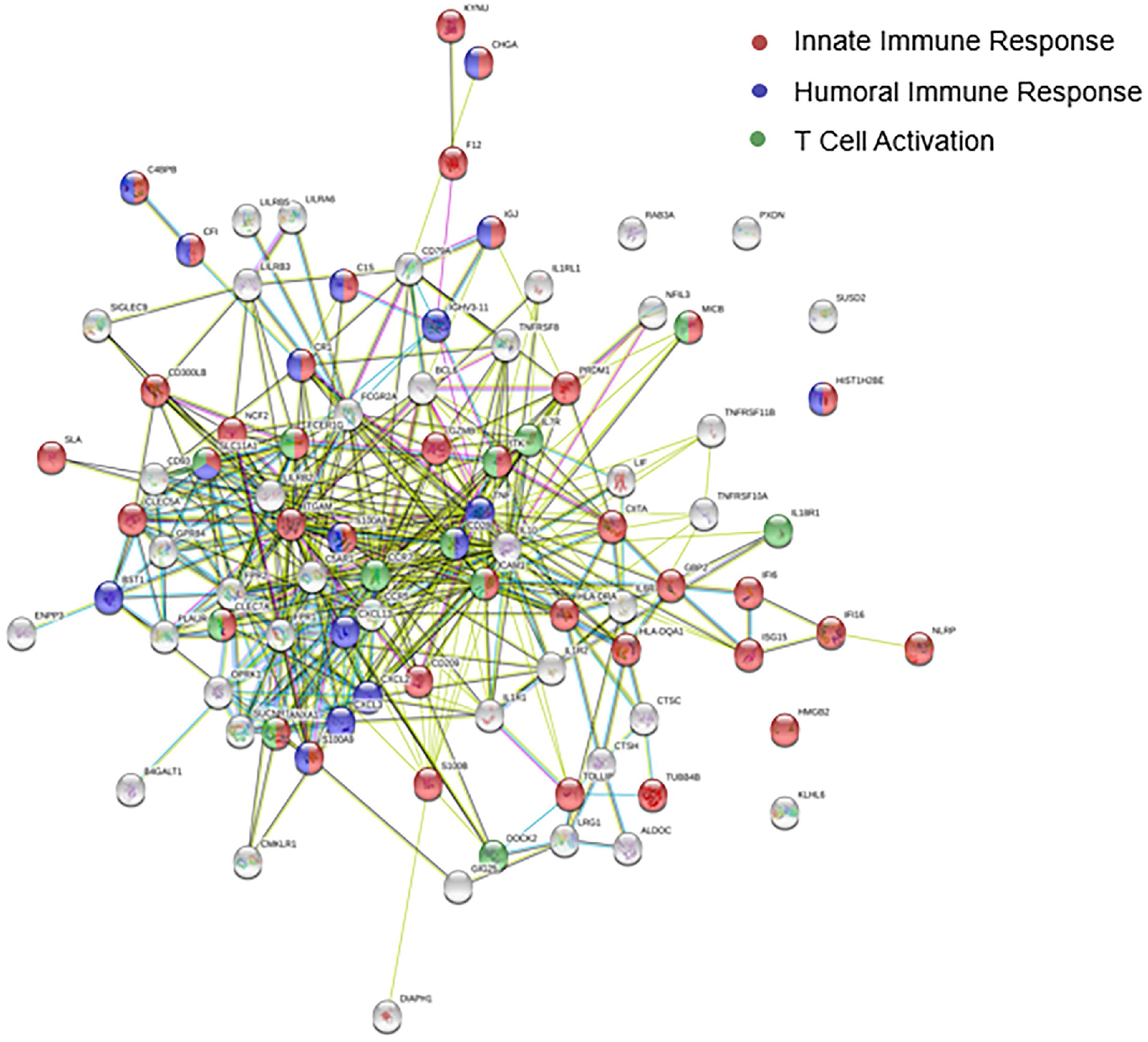
Interaction network of immune responses: 89 genes from our DEG list were registered as immune response related genes (GO:0006955). The immune response genes were subsequently reentered into STRING to identify the nature of the inflammatory reactions and to determine possible protein network interactions. Gene changes related to both adaptive and innate immune responses were recognized in our gene list. 38 genes (red) are considered part of an innate immune response (GO:0045087), while 17 genes (blue) are associated with a humoral immune response (GO:0006959) and 12 genes (green) are involved in T cell activation (GO:0042110).

**Supplemental Table 1**

RNA quality table with RNA integrity number, Phred quality score, read count, and % uniquely mapped reads.

**Supplemental Table 2**

Significant genes (adjusted p-value cutoff of 0.05 by Benjamini Hochberg’s False Discovery Rate).

**Supplemental Table 3**

A list of all genes, including base mean, log2 fold change, and padj values.

**Supplemental Table 4**

Normalized data (DESEQ2 normalized counts)

**Supplemental Figure 5**

Table of 71 dysregulated immunoglobulin genes including IGHG1-4, IGHA1-2, and IGHM.

**Supplemental Table 6**

Further gene enrichment was conducted using ToppGene Suite (https://toppgene.cchmc.org). In the DisGeNET database of gene-disease associations, 79 dysregulated genes were listed as involved in pain (C0030193).

Piñero J, Ramírez-Anguita JM, Saüch-Pitarch J, Ronzano F, Centeno E, Sanz F, Furlong LI. The DisGeNET knowledge platform for disease genomics: 2019 update. Nucleic Acids Res. 2020 Jan 8;48(D1):D845-D855. doi: 10.1093/nar/gkz1021.

**Supplemental Table 7**

Our list of DEGs was cross-referenced with mouse homologs registered in the Pain Genes Database (http://www.jbldesign.com/jmogil/enter.html), which documents the pain testing performed genetically engineered null mice from the literature (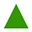 = mutant more sensitive, 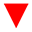 = mutant less sensitive, 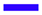 = no difference, 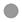 = not tested, 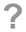 = contradictory data)

LaCroix-Fralish, M.L., Ledoux, J.B. and Mogil, J.S. The Pain Genes Database: an interactive web browser of pain-related transgenic knockout studies. Pain, 131:3.e1-3.e4, 2007.

**Supplemental Table 8**

Since axonal regeneration may contribute to pain in diabetic neuropathy patients, The STRING Database was used to identify upregulated genes that could be associated with nervous system development (GO:0007399).

