## Supplemental data for "Transcriptomic Analysis of Human Sensory Neurons in Painful Diabetic Neuropathy Reveals Inflammation and Neuronal Loss"

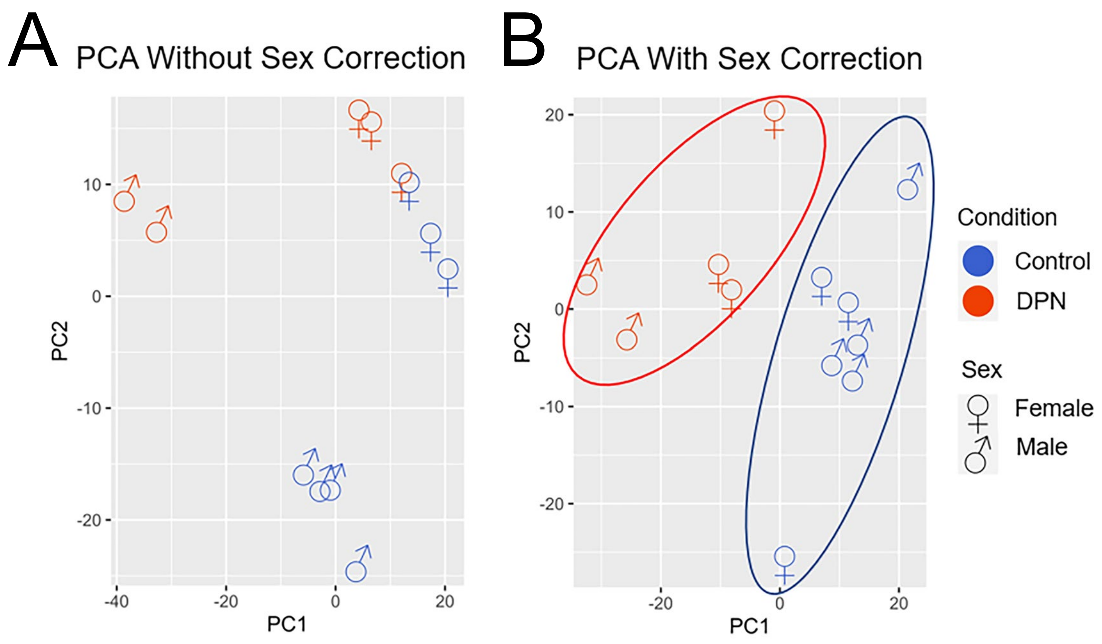


**Supplemental Figure 1**

Principal component analysis (PCA) of human DPN transcriptome data: PCA plot A) before and B) after including sex as a covariate as part of the DESEQ2 analysis. PC1 is the first principal component direction where the most variance is occurring, and PC2 is the second most one that is orthogonal to PC1. With sex as a covariate, the DPN donors and the controls then separate into two independent groups.

A

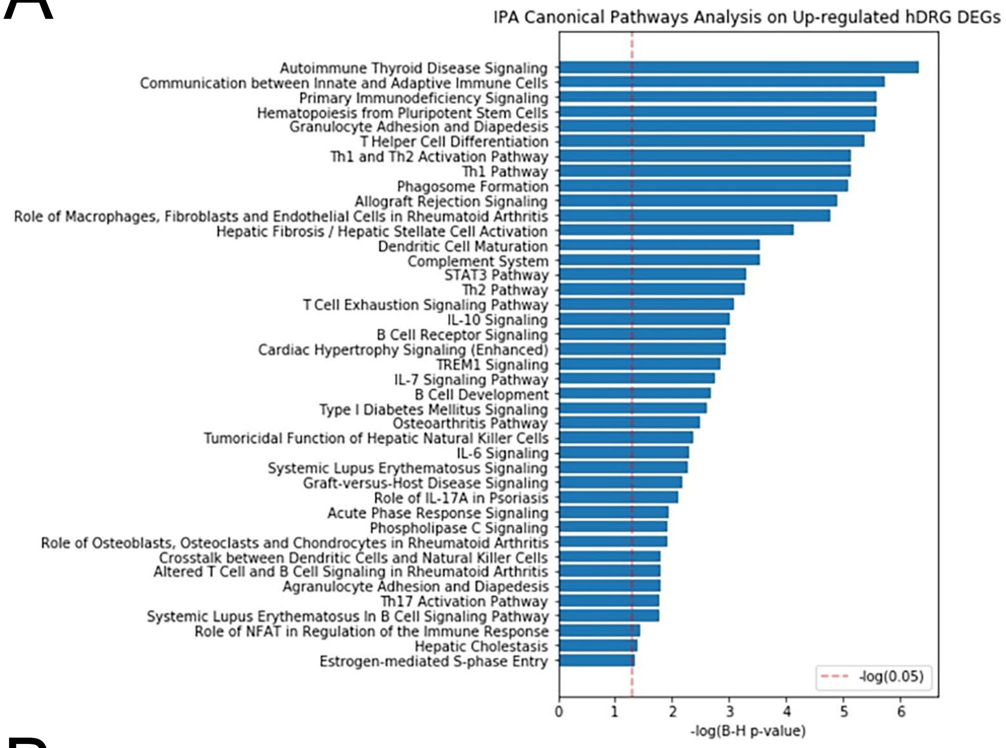


B

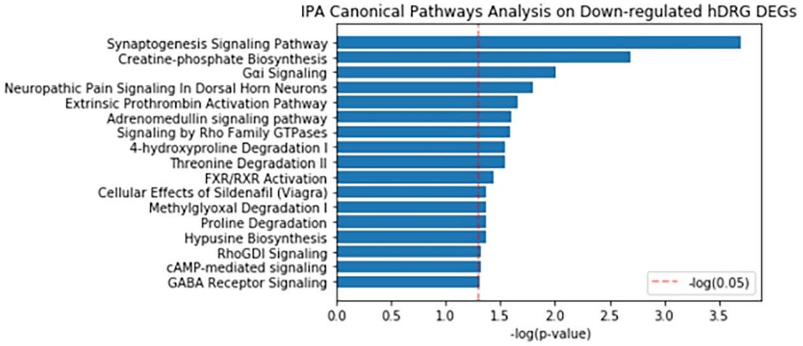


**Supplemental Figure 2**

Ingenuity pathway analysis (IPA) of transcriptomic data: The IPA report using all 844 dysregulated genes predominantly centered on the immunological functions occurring in the DRG of the DPN individuals. Thereby, further assessment of the DEGs was separated between A) the upregulated largely inflammatory gene responses and B) the downregulated gene expression changes, where synaptogenesis was appears to be affected by decreases in gene expression.

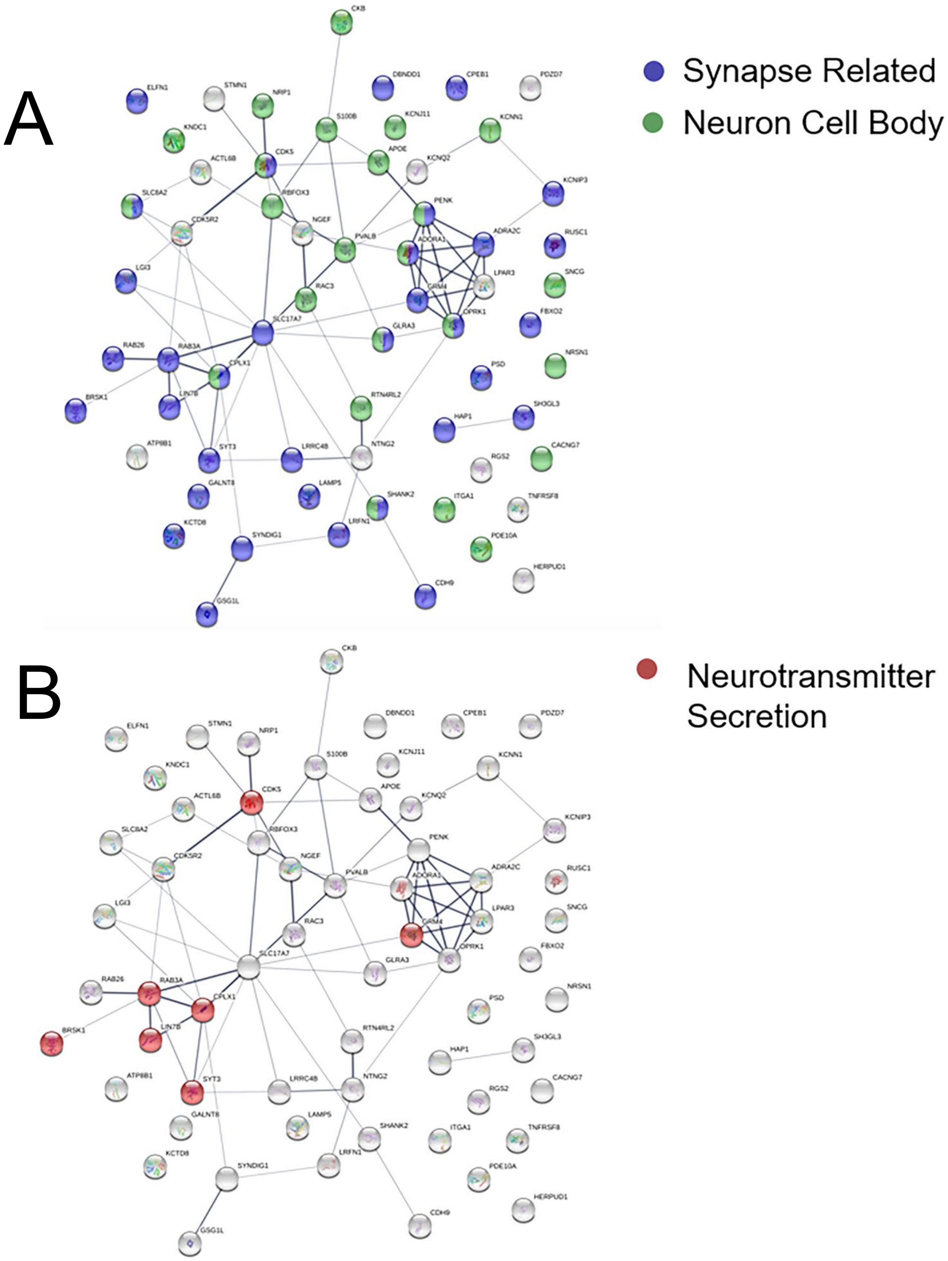


**Supplemental Figure 3**

Interaction network of neuronally related genes: 62 genes out of our DEG list were considered to perform as a cellular component of a neuron (GO:0097458). Most genes were downregulated (n=51) while a few were upregulated (n=11) in the DPN donors. To further determine how these dysregulated genes might impact neuronal function, all 62 gene were separately evaluated using STRING (https://string-db.org/) for additional enrichment analysis. A) About 66% of the neuronal genes (n=41) were synaptically related (blue - GO:0045202), while 39% (n=24) were associated with the neuron cell body (green - GO:0043025). B). In terms of biological function, a few genes in red were qualified as being associated with neurotransmitter secretion (GO:0007269).

Kuleshov MV, Jones MR, Rouillard AD, Fernandez NF, Duan Q, Wang Z, Koplev S, Jenkins SL, Jagodnik KM, Lachmann A, McDermott MG, Monteiro CD, Gundersen GW, Ma'ayan A. Enrichr: a comprehensive gene set enrichment analysis web server 2016 update. Nucleic Acids Research. 2016; gkw377

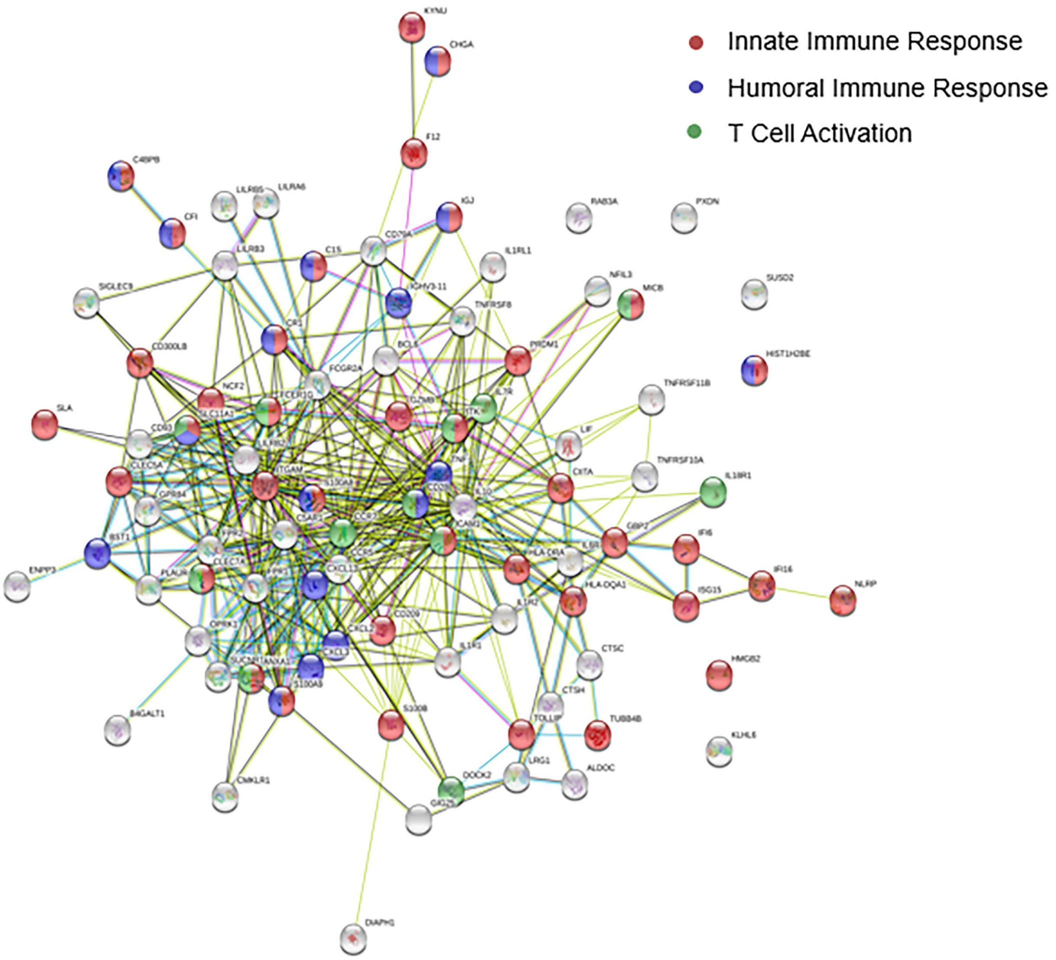


**Supplemental Figure 4**

Interaction network of immune responses: 89 genes from our DEG list were registered as immune response related genes (GO:0006955). The immune response genes were subsequently reentered into STRING to identify the nature of the inflammatory reactions and to determine possible protein network interactions. Gene changes related to both adaptive and innate immune responses were recognized in our gene list. 38 genes (red) are considered part of an innate immune response (GO:0045087), while 17 genes (blue) are associated with a humoral immune response (GO:0006959) and 12 genes (green) are involved in T cell activation (GO:0042110).

**Supplemental Table 1**
RNA quality table with RNA integrity number, Phred quality score, read count, and % uniquely mapped reads.

**Supplemental Table 2**
Significant genes (adjusted p-value cutoff of 0.05 by Benjamini Hochberg's False Discovery Rate).

**Supplemental Table 3**
A list of all genes, including base mean, log2 fold change, and padj values.

**Supplemental Table 4**
Normalized data (DESEQ2 normalized counts)

**Supplemental Figure 5**
Table of 71 dysregulated immunoglobulin genes including IGHG1-4, IGHA1-2, and IGHM.

**Supplemental Table 6**
Further gene enrichment was conducted using ToppGene Suite (https://toppgene.cchmc.org). In the DisGeNET database of gene-disease associations, 79 dysregulated genes were listed as involved in pain (C0030193).

Piñero J, Ramírez-Anguita JM, Saüch-Pitarch J, Ronzano F, Centeno E, Sanz F, Furlong LI. The DisGeNET knowledge platform for disease genomics: 2019 update. Nucleic Acids Res. 2020 Jan 8;48(D1):D845-D855. doi: 10.1093/nar/gkz1021.

**Supplemental Table 7**
Our list of DEGs was cross-referenced with mouse homologs registered in the Pain Genes Database (http://www.jbldesign.com/jmogil/enter.html), which documents the pain testing performed genetically engineered null mice from the literature (▲ = mutant more sensitive, ▼ = mutant less sensitive, ▬ = no difference, ● = not tested, ? = contradictory data)

LaCroix-Fralish, M.L., Ledoux, J.B. and Mogil, J.S. The Pain Genes Database: an interactive web browser of pain-related transgenic knockout studies. Pain, 131:3.e1-3.e4, 2007.

**Supplemental Table 8**
Since axonal regeneration may contribute to pain in diabetic neuropathy patients, The STRING Database was used to identify upregulated genes that could be associated with nervous system development (GO:0007399).
